# Deterministic DNA-first discovery reveals recurrent genomic architectures

**DOI:** 10.64898/2026.11.01.763506

**Authors:** Guoqing Hu, Hao Wang, Stephen S.-T. Yau

## Abstract

Genome mining typically begins with known genes or protein families, which can make systems defined primarily by genomic organization difficult to detect. We developed a DNA-first framework combining asymmetric-ovariance natural-vector representations with generic positional recurrence, joining separately calibrated signals through genomic coordinates. In benchmarks of array-associated reverse transcriptase (ART) loci, the target-free mathematical arm localized seven of eight predeclared loci, including all seven with exact-array coordinates. Dual-arm signals occurred in none of three non-array RT controls and in one of five UG27 systems, indicating selectivity within these small control sets. A separate profile-guided search of 6.12 Gb identified five candidate array-associated RT-like systems. In an independent 54.38-Gb evaluation of this profile-guided arm, structural compatibility was 24/32 under a historical-rule mirror, 12/32 after novelty-only selection, and 0/17 for architecture-selected complete ORFs; the two 32-member panels shared six exact sequences. Together, these results suggest that mathematical features of raw DNA can complement protein-directed searches in nominating genomic architectures for experimental RNA-to-DNA studies.

## Introduction

Genome analysis commonly begins with genes, protein families and functional annotations. Protein-family resources, functional-annotation tools and genome-mining methods identify different classes of known sequence features [1–6]. Related searches for microbial defence systems use conserved domains, curated system definitions or genomic neighbourhoods to identify and classify candidate loci [7–10]. These approaches often require investigators to specify the molecular class of interest before examining the surrounding DNA. Genomic co-localization and alternative functional screens have also revealed previously unrecognized antiphage systems [11, 12]. These strategies are effective when the target class is known, but unusual DNA organization might also provide a way to identify candidate systems before their protein families or mechanisms are established.

Prokaryotic reverse transcriptases participate in diverse mobile-element and defense-associated systems [13–15]. Retrons, for example, combine an RT with noncoding RNA and, in some systems, an effector protein [16, 17]; their antiphage activity and phage countermeasures have been examined experimentally [18]. The diversity of bacterial antiphage mechanisms provides a broader context for system-level discovery [19, 20]. Array-associated reverse transcriptases (ARTs) provide a test case. The reported agentic ART campaign began from a massive protein-centric search and an autonomous agent later noticed an atypical repeat array by reading local DNA around an unusual reverse transcriptase (RT) [21]. The MarsHill RT had been described previously, but the repeat–RT–partner archi-tecture was recognized only when the locus was examined as a system [22]. This example illustrates the difference between identifying a known sequence feature and recognizing its surrounding genomic architecture.

Sequence representation methods include alignment-free mathematical approaches and pretrained DNA language models [23–27]. Such language models have been studied as representations of genomic sequence context [28, 29]. Here we examine whether mathematical features of raw DNA can nominate genomic regions for further study without first defining a target protein family. We therefore separate candidate nomination from biological interpretation. Arm M begins from nucleotide sequence alone and combines the asymmetric-covariance natural vector (ACNV), which measures positional organization, with generic sequence recurrence. A second, orthogonal arm begins from raw DNA but enters protein space through six-frame translation and frozen RT profiles. The two arms operate independently: one identifies unusual genomic organization without specifying a target protein class, whereas the other identifies RT-profile-related proteins from translated raw DNA without relying on gene annotation. Biological interpretation is performed only after their respective discovery outputs are frozen (Fig. 1).

**Figure 1:**
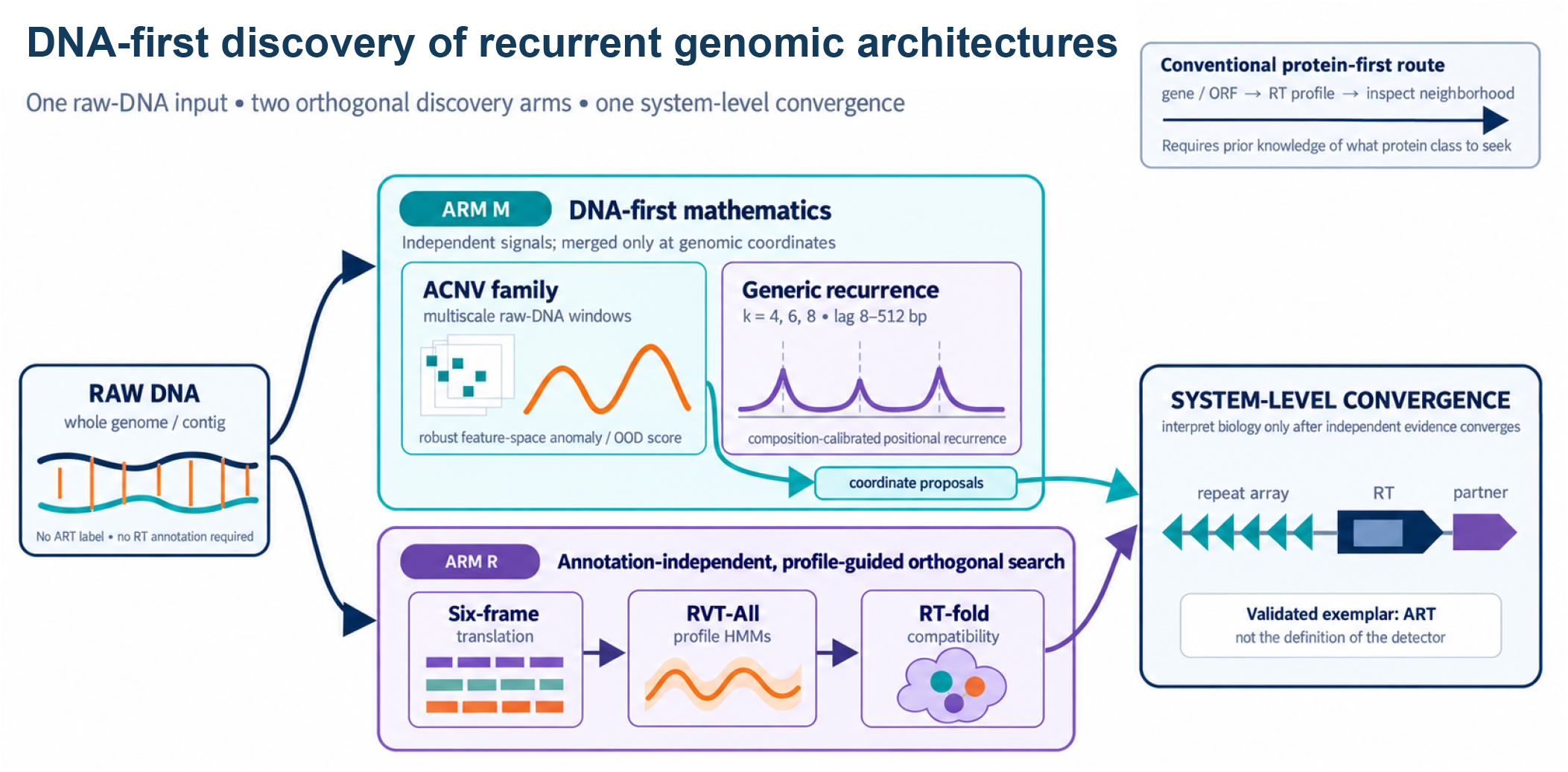
DNA-first discovery of recurrent genomic architectures. Raw DNA is exposed to two orthogonal analyses before system-level interpretation. In Arm M, multiscale ACNV and generic positional recurrence are scored independently and meet through genomic-coordinate proposals; no RT coordinate, ART label, repeat motif or partner annotation enters the DNA-scoring layer. Arm R is annotation-independent and profile-guided: six-frame peptides are screened by frozen RVT-All profiles and assessed against a fixed clean-RT structural reference set. The inset shows a conventional target-aware RT-first route. Each arm has its own frozen outputs; the convergence panel depicts subsequent interpretation of repeat, RT and partner evidence. ART is an established exemplar of this broader discovery object. The five newly prioritized systems remain computational candidates pending causal experiments.

## Results

### Independent mathematical views nominate genomic coordinates before biological in-terpretation

The natural-vector family characterizes biological sequences through composition and positional summaries, including k-mer extensions [30–32]; the energy entropy vector is a related, distinct fixed-dimensional representation [33]. ACNV was introduced as a fixed-dimensional representation that preserves direction-sensitive positional covariance in biological sequences [34]. Here it is used as a representation rather than a supervised ART decision rule. For each feature *j* within a fixed sequence, window-length and strand stratum, we robustly standardize

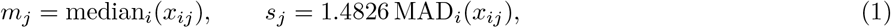

using a mean-absolute-deviation fallback when the robust scale is at most 10^*−*12^ and excluding constant features. We then define

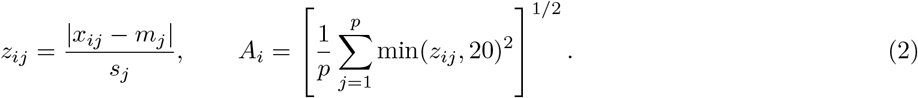

Scores are percentile-ranked separately within each window length and strand of the scored sequence. The larger strand percentile is retained and re-ranked within length. The frozen ACNV family contains ACNV92 and separate 4-, 5- and 6-mer specialists; their maximum surprisal at each window is again ranked among windows of the same length (Supplementary Fig. S1). Throughout, *q* denotes an empirical upper-tail fraction used for nomination, not a false-discovery-rate estimate. The implementation and tie rules are given in the Supplement.

Repeat and motif discovery has a substantial methodological literature, including tandem-repeat finding, CRISPR-array recognition and motif analysis [35–37]. Generic recurrence is computed independently. For *k ∈* {4, 6, 8 and} lags *d* = 8, …, 512 bp, exact *k*-mer reappearance is normalized using the local-composition expectation

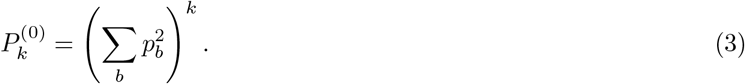

No ART-specific lag or motif is supplied. ACNV therefore asks whether positional organization is unusual, while recurrence asks whether short sequence patterns reappear at unexpected separations. Windows meeting the frozen empirical-tail threshold in either view are merged by genomic-coordinate overlap.

L0050 provides a concrete worked example (Fig. 2). After score freeze, the published upstream region reconstructed to 153,152–156,052 bp. The frozen dual-arm region spanned 153,088–157,696 bp and ranked 1/84. The strongest ACNV window was 154,240–154,368 bp (family *q* = 2.80 *×* 10^*−*4^; ACNV6MER_BLOCK), while the strongest recurrence window was 155,136–155,392 bp (*k* = 8, *q* = 1.40 *×* 10^*−*4^). The full frozen score matrix independently reproduced all 137 selected proposals and 84 merged regions. Recalculation from the 256-bp recurrence-peak DNA reproduced *k* = 8, a best lag of 154 bp and 15 exact matches among 95 eligible pairs. This directly supports a 154-bp recurrence scale. Earlier post-score descriptions also report 14 repeat copies, median start-to-start spacing of 154 bp and the 16-nt consensus CATGTGTATCGCATGT; the original copy-level alignment is unavailable, so this descriptive reconstruction remains distinct from the independently replayed score and region results.

**Figure 2:**
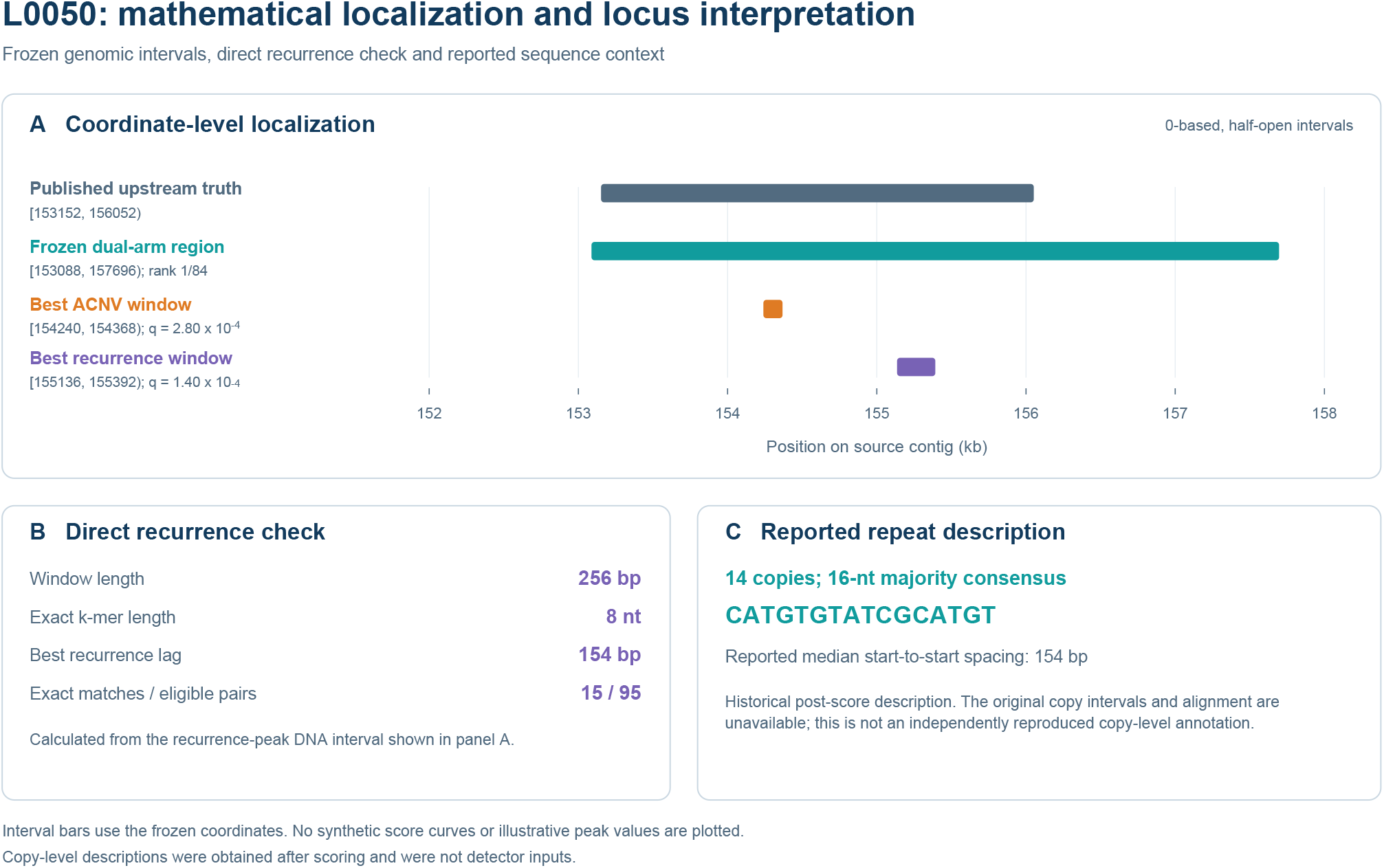
Worked example: mathematical localization and locus interpretation of L0050. **A**, Coordinate bars show the published upstream truth interval [153152, 156052), the frozen dual-arm region and the best ACNV and recurrence windows on a shared genomic axis. **B**, The direct recurrence check reports the peak-window length, *k*, best lag and exact-match count. **C**, The historical post-score repeat description is reported separately because the original copy intervals and alignment are unavailable. The frozen merged dual-arm region is [153088, 157696), rank 1/84; the best ACNV window is [154240, 154368) with family *q* = 2.80 *×* 10^−4^; and the best recurrence window is [155136, 155392) with *k* = 8 and *q* = 1.40 *×* 10^−4^. These sourced intervals are 0-based and half-open. The independently recalculated best lag is 154 bp. Earlier summaries report 14 repeat copies, median 154-bp spacing and the 16-nt consensus CATGTGTATCGCATGT; the copy-level provenance limitation is detailed in the Supplement. No synthetic score curves or illustrative peak values are plotted. Here *q* is an empirical within-contig, within-length upper-tail fraction.

### Known ART architectures are prioritized without using ART or RT labels

Historical development and validation were kept separate. MarsHill is explicitly retained as the development genome, including its original stringent blind near-miss, rather than rewritten as an untouched success. After the region-integration and ranking rules were fixed, the final candidate tables placed the known loci at ranks 1/84 for L0050, 2/84 for MarsHill, 2/82 for Madawaska, 3/78 for SA1 and 4/80 for Machias. Earlier post-score descriptions describe repeat organization upstream of the RT, including a reported median spacing of 154 bp in L0050 that agrees with the independently recalculated recurrence lag (Fig. 3). These motif and spacing descriptions were unblinding products and were never detector inputs; the Supplement distinguishes available sequence-level evidence from historical copy and anchor records that could not be independently replayed.

**Figure 3:**
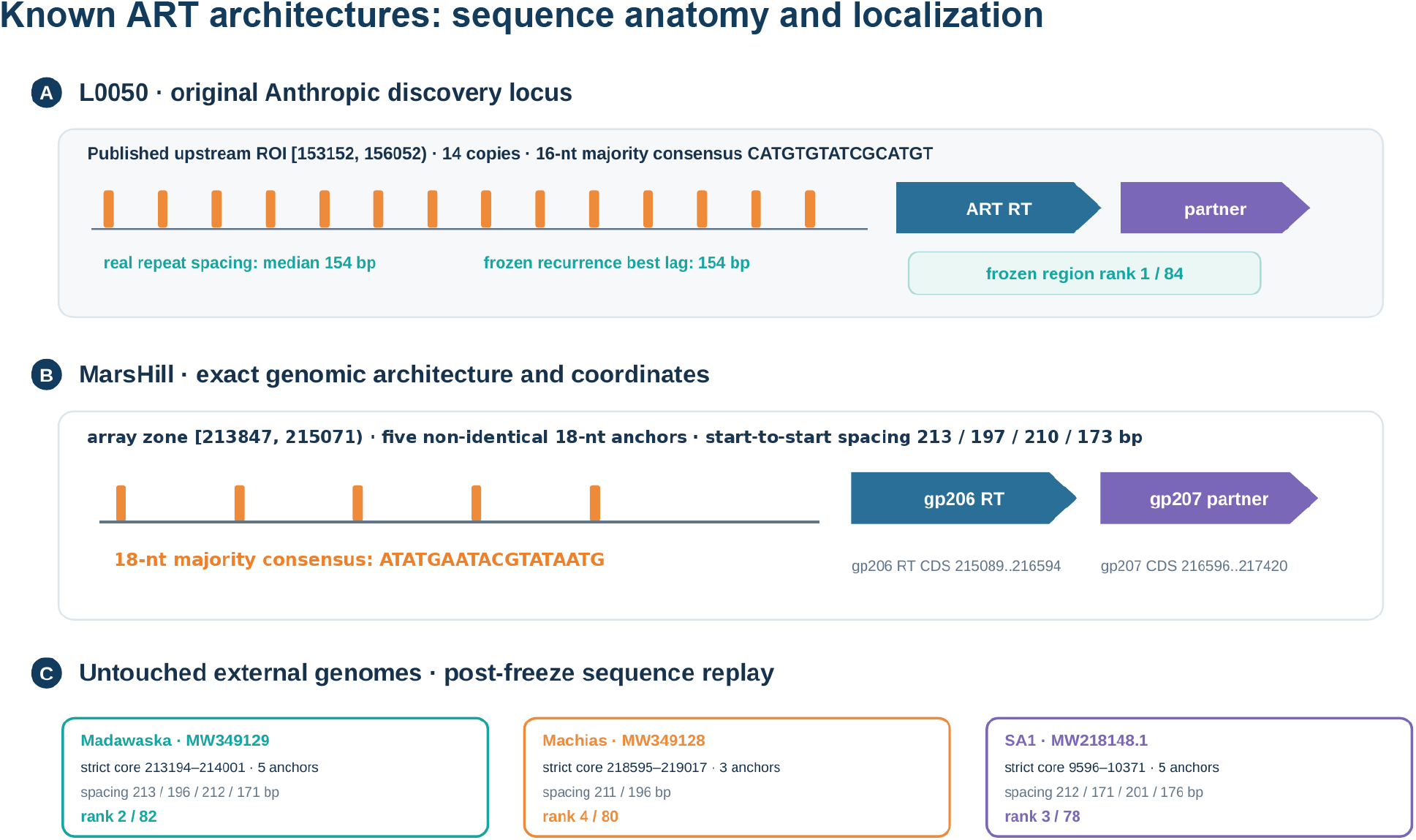
Sequence anatomy and mathematical localization of benchmark ART loci. **A**, L0050 is a locked retrospective case with a reported 14-copy reconstruction and median spacing of 154 bp, independently recalculated best recurrence lag of 154 bp, and rank 1/84. **B**, MarsHill retains its development-locus annotation and five reported post-freeze replay anchors upstream of gp206 RT and gp207 partner. **C**, Madawaska, Machias and SA1 are external cases, with strict anchor intervals, spacings and final clean ranks. Exact/strict anchor counts are distinct from full reconstructed copy counts. Repeat and protein glyphs are schematic and are not drawn to nucleotide scale. Source-verified coordinate conventions and evaluation roles are provided in Supplementary Data; the original MarsHill CDS notation is retained. The MarsHill motif is a reported descriptive consensus, distinct from the separately reported 18-nt anchors. The reference-DNA audit and remaining copy/anchor provenance limitations are detailed in the Supplement.

### A frozen conventional benchmark separates target-aware localization from target-free discovery

Target-free mathematical screening and RT-directed searches use different prior information and address different localization tasks. We therefore predeclared eight ART-related test cases (Madawaska, Machias, SA1, LY01, S6, PALS_2, UFV_DC4 and PB50), froze all candidate boards before truth unblinding, and compared the mathematical arm with frozen RT-first searches, including a Prodigal-to-RVT profile route [38, 39]. Protein-first candidates were RT-centered *±*10-kb neighborhoods; adding recurrence used the same frozen recurrence evidence as Arm M.

Both predeclared protein-first primary comparators—Prodigal–RVT plus frozen recurrence and six-frame–RVT plus frozen recurrence—localized 8/8 cases (6/8 at rank 1; MRR 0.875 for each). The corresponding secondary RVT-only comparators each localized 8/8 at rank 1. Arm M recovered 7/8 (MRR 0.406) and retained PB50 as the prespecified miss. On the seven cases with exact array truth, however, Arm M recovered 7/7, placing 6/7 within the top three and 7/7 within the top five genome-wide regions; all three strictly external array loci were recovered without retuning. Coordinate-level convergence was essential for prioritization: on these seven exact-array cases, full ACNV+recurrence achieved MRR 0.464, compared with 0.084 for ACNV alone and 0.151 for recurrence alone (Table 1; Supplementary Table S3). Thus the contribution of recurrence is not that it universally improves an RT HMM: when appended to either target-aware RT-first baseline it reduced rank performance (MRR 1.000 to 0.875 on the eight-case panel). Its value emerges as an independent mathematical view within a biology-blind whole-genome search. Independent reevaluation of the frozen canonical boards reproduced all 77 archived case–method records, including both primary protein-first contrasts.

**Table 1:**
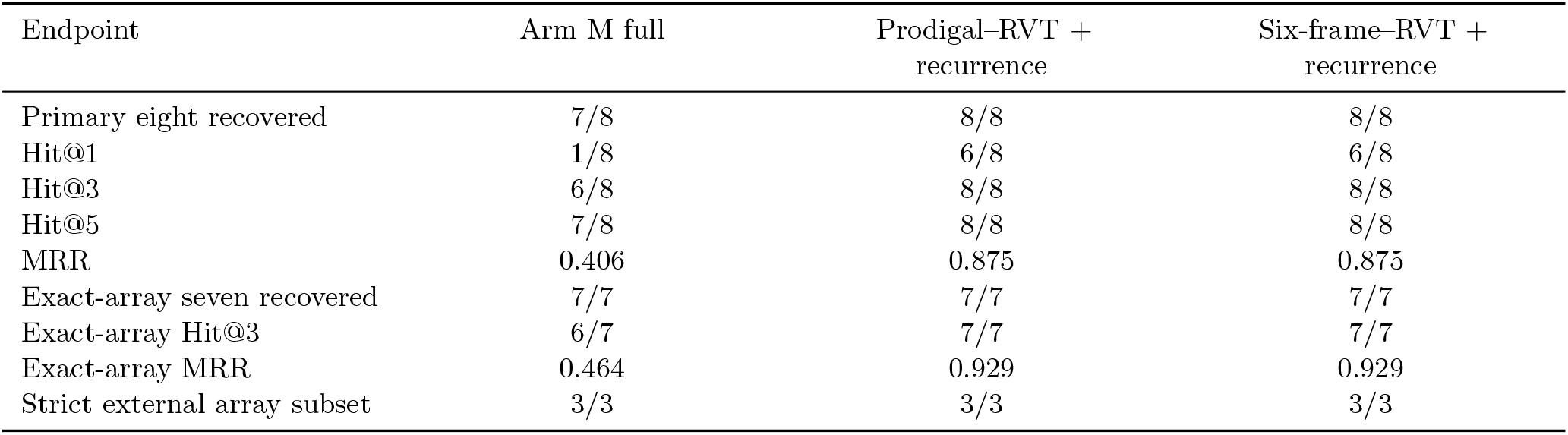
Frozen conventional-method benchmark: three predeclared primary contrasts. The primary panel retains all eight cases, including the Arm-M miss PB50. Exact-array and strictly external subsets are reported separately. The two predeclared secondary RVT-only routes each have eight-case Hit@1 8/8 and MRR 1.000; full method details and case-level results remain in Supplementary Table S3 and Supplementary Information, Section 7.

When RT identity is specified in advance, protein-first searches rank these known loci more effectively. Arm M addresses a different setting: genome-wide nomination without assuming that RT, or another protein family, is the relevant target. Candidate burden is therefore reported as the number of regions requiring review, not as a false-positive rate.

### Specificity is architectural: non-array RTs and UG27 define two independent boundaries

Three mechanistically established non-array RT systems—Sen2, Ec78/Eco7 and DRT2—provide a stringent control against interpreting the method as an RT detector. Their RT loci can be ACNV-anomalous, but none satisfied the independent recurrence requirement; none of the three met the dual-arm criterion (0/3). Their mathematical-union ranks (186/1477, 350/1493 and 188/1636, respectively) further show that the result is not simply failure to register local sequence anomaly.

UG27 provides a harder near-neighbor. These experimentally studied systems couple an RT to structured ncRNA arrays whose primary sequences can be diverse [40]. Crucially, the UG27 experiment was scored genome-wide before reference locus annotations were accessed. Across five native assemblies (15 contigs, approximately 20.66 Mb), the frozen score board contained 6,212 clean ACNV/recurrence union regions and 303 clean dual-arm regions. Cassette-level unblinding showed direct ACNV overlap in 5/5 systems but dual-arm overlap in only 1/5.

We then defined the ncRNA reference intervals independently of the detector by mapping the published, machine-readable example-locus annotations. Thirty-one annotated ncRNA units were extracted from the five example loci; 28 mapped uniquely and exactly inside the independently frozen experimental cassettes (8, 6, 6, 6 and 2 units per system). The three excluded Blautia units lay outside the experimental cassette and were not included in the cassette-level comparison. Re-evaluation against these mapped ncRNA units reproduced the same boundary: only *Alistipes putredinis* was exact-unit dual-arm positive, with 3/8 units directly overlapping one frozen dual-arm region. Each of the other four systems nevertheless retained direct exact-unit ACNV-only support in two units (Supplementary Table S1). No score, region board, threshold, lag range or nearby-distance rule was recomputed after the reference intervals were defined.

The descriptive pattern is 5/5 dual-arm across the five illustrative ART loci (one development, one locked retrospective and three external cases); the UG27 near-neighbor is 1/5 exact-array dual-arm despite exact-unit ACNV support in 5/5; and mature non-array RT controls are 0/3 dual-arm. These small, ascertained panels define an empirical boundary rather than population specificity or an ART-specific biochemical signature: RT presence and an ncRNA array alone do not ensure dual-arm nomination.

A broader analysis of 919 genomes yielded 3,674 mathematically nominated regions and 602 repeat-linked loci. Frozen E1 source memberships (1,166 edges), rules and D1/D4R1 replay were recovered and reproduced 66 replicated families (324 loci; 250-bp criterion). The prespecified 1,000-bp sensitivity analysis yielded 74 families (388 loci). Recovered E2A membership and exact-primary/FAM50 criteria reproduced 12 high-specificity families (119 loci) within the frozen 999-candidate audit universe. These counts are computational, not functional or global-novelty claims. The archived N2 native-DNA audit reports completed repeat-group, copy and spacer coordinate checks for all 602 loci, covering 16,527 groups, 64,988 copies and 48,461 spacers with no reported discrepancies. This coordinate verification is distinct from a fresh 919-genome scan or biochemical validation (Supplementary Table S8, P02/P03/P10; Supplementary Information, Section 21.3).

### An orthogonal 6.12-Gb raw-DNA search converges on five computational ART-like systems

Arm R tests a different discovery route. A 6.1216-Gb external raw-DNA universe was translated in all six reading frames into maximal stop-to-stop segments, avoiding dependence on gene calling. Approximately 65.3 million peptide segments were screened against a frozen 45-profile RT collection derived from myRT [39] using HMMER [41]. The screen produced 10,598 formal RVT-profile loci, which were deduplicated and clustered with MMseqs2 [42] into 4,912 operational families. Thirty-two priority families were frozen under a fixed structural-compute budget. Predicted structures were obtained for 30 representatives, while two remained technically unresolved; 28 of the 30 attained at least suggestive structural compatibility with the fixed clean-RT reference set using Boltz-2 and Foldseek [43, 44].

System-level reconstruction of the structural survivors examined upstream repeat architecture and recurrent down-stream partner candidates without using existing annotation as a hard discovery filter. Five systems remained high priority after this reconstruction. One prioritized family combines strong RT-fold support and three near-context clusters; its selected native architecture contains an ATGGAGTA repeat core with 157/157-bp inter-repeat spacers (165/165-bp start-to-start intervals) and a conserved 125-aa same-strand downstream partner beginning 13 bp after the RT terminal stop. The five labels denote computational convergence of architecture and RT evidence, not biochemical confirmation.

### An independently frozen 54-Gb universe separates transfer, novelty selection and architecture-first discovery

The approximately 54-Gb experiment evaluates the profile-guided search on an independent source universe; it is not a 54-Gb genome-wide validation of the biology-blind mathematical arm. The initial discovery universe contained 6,121,620,524 bp from 131 source accessions. The independent universe contained 54,377,819,157 bp from 254 newly selected source accessions, with all 131 original accessions excluded before freeze. Acquisition retained the same four broad source strata—viral/virome, environmental metagenome, host-associated metagenome and extreme/underrepresented sources—but the accession-level membership, taxonomic composition and contig realization were independent. Six-frame translation, formal RVT-profile rules, family compression, structural reference set and query-level structural thresholds were unchanged.

The independent scan produced 88,487 formal RT-like instances, 85,503 exact-AA-unique sequences and 26,396 native operational families, compared with 10,598, 10,329 and 4,912, respectively, in the initial universe. Formal RT-like density declined only from approximately 1,731 to 1,627 instances per Gb (about 6%), whereas operational-family density declined from approximately 802 to 485 families per Gb, showing lower family density in this sampled universe. Because source composition differs, this density comparison alone does not establish discovery saturation.

We then kept three 54-Gb routes explicitly separate (Fig. 5). First, a *strict mirror* replayed the historical four-axis rule—replication, maximum amino-acid length, formal model coverage and motif-topology, with deterministic backfill—directly over the 26,396 native operational families, without a combined old–delta novelty filter. This yielded 32 representatives: 18 strong, six suggestive, seven weak and one no-hit, or 24/32 (75.0%) at least suggestive. Frozen execution manifests record 26 new structure predictions, six exact-amino-acid structure reuses and 32 completed Foldseek searches. Independent reclassification of all 901 archived strict-mirror hits reproduced the 18/6/7/1 distribution, and all 32 predicted-structure polymer sequences matched the frozen inputs. These classifications describe protein-level structural compatibility; they do not establish 24 complete ART systems or biochemical function. No additional manual structural review was performed. Second, joint old–delta clustering identified 24,810 delta-only combined families after exclusion of mixed historical–delta families. Applying the same frozen four-axis selector to this novelty-filtered family space produced the *novelty-only protein route*: three strong, nine suggestive, 18 weak and two no-hit, or 12/32 (37.5%) at least suggestive. The strict-mirror and novelty-only panels shared only six exact amino-acid sequences out of 32, demonstrating that novelty-oriented filtering substantially changes which proteins reach structural evaluation. The original 28/30 versus 12/32 contrast must therefore not be interpreted as a like-for-like replication rate; the comparison under the same selection and grading rules is 28/30 historical versus 24/32 strict mirror, with attenuation but substantial structural transfer. The two technically unresolved historical representatives remain outside the 30-member denominator and are not counted as structural negatives.

**Figure 4:**
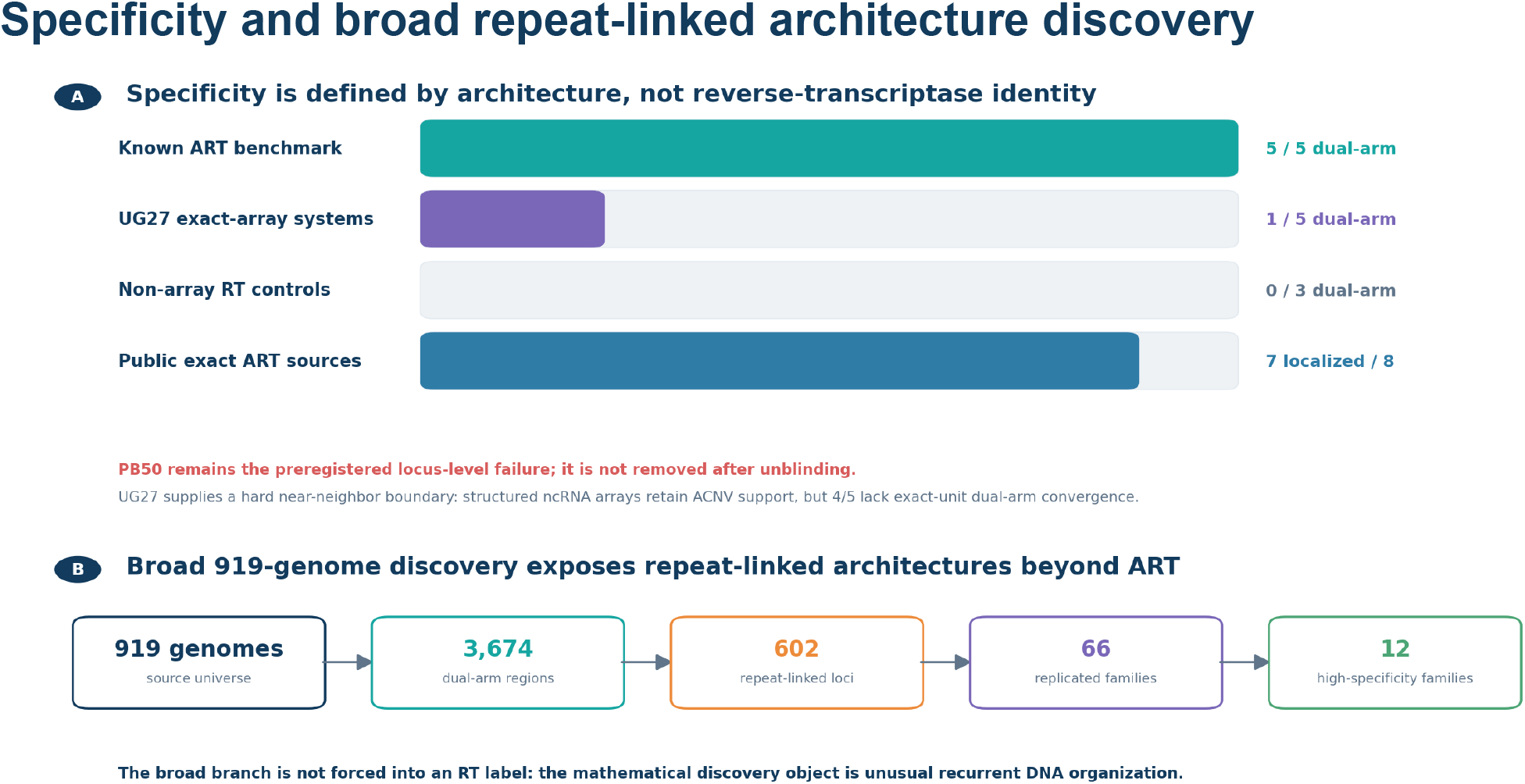
Specificity and broad repeat-linked architecture discovery. **A**, The panel summarizes 5/5 dual-arm overlap among five illustrative ART loci, 1/5 among UG27 exact-array systems, 0/3 among non-array RT controls, and localization of seven of eight public ART-related source loci. The five illustrative cases comprise one development, one locked retrospective and three external cases. These are distinct endpoints from small ascertained sets and do not estimate population specificity. PB50 remains a locus-level miss in the separate eight-case benchmark; the primary comparator definitions and available results are retained in Supplementary Table 1. **B**, The 919-genome branch contains 3,674 mathematical regions and 602 repeat-linked loci. Its manifest matches the frozen checksum, and the region records and 602-locus selection were independently checked. The displayed 66 replicated-family and 12 high-specificity-family labels come from an earlier figure source-data snapshot; original calculation, membership and specificity-criterion records were not recovered. These two values are unverified historical annotations, not independently supported family-level results.

**Figure 5:**
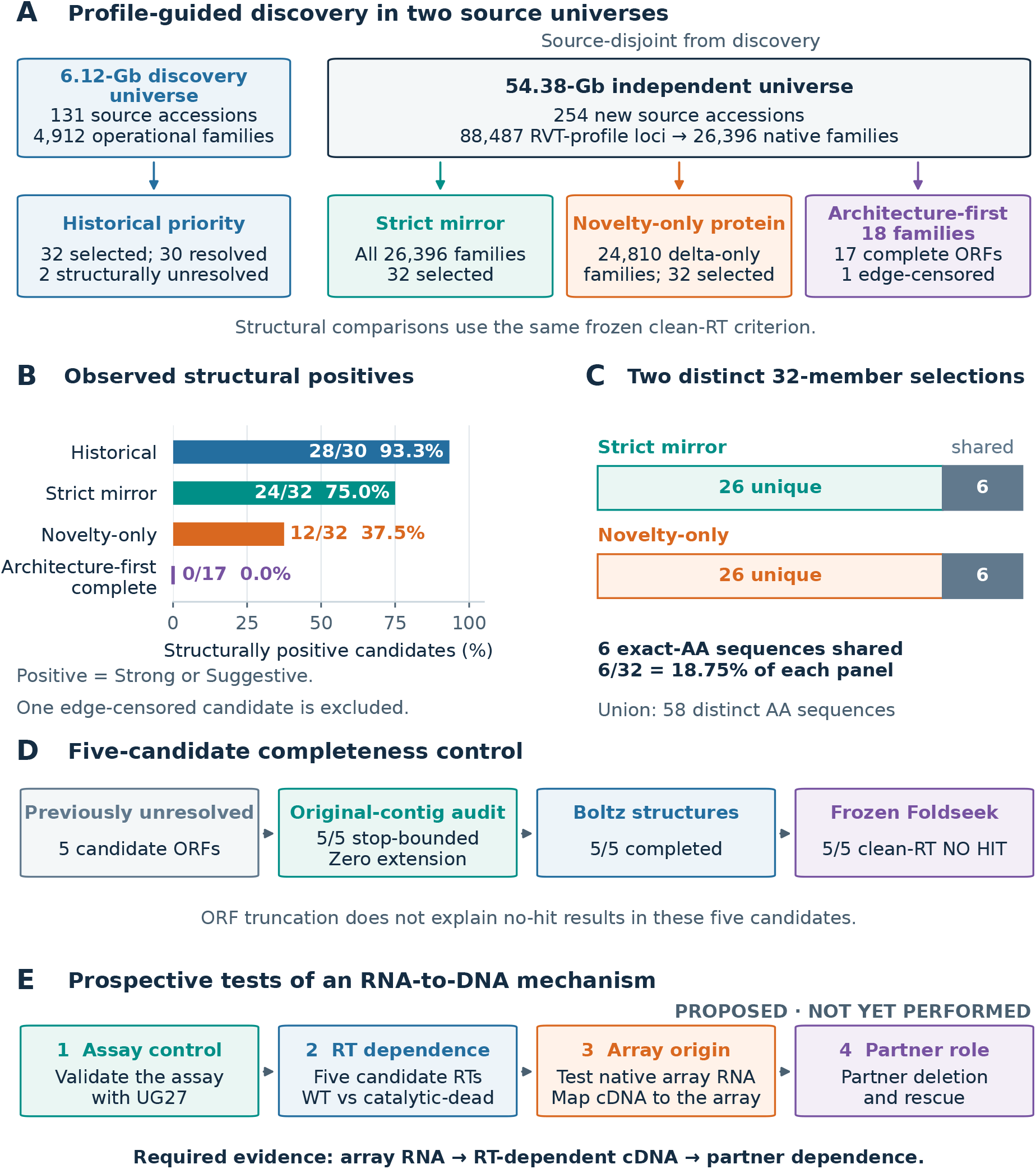
Candidate selection and structural outcomes in independent source universes. **A**, All displayed routes begin from RVT-profile candidates. Architecture-first selection prioritizes genomic recurrence before structural ranking and remains profile-conditioned. **B**, Strong or suggestive clean-RT compatibility is observed in 28/30 historical resolved, 24/32 strict-mirror, 12/32 novelty-only and 0/17 complete architecture-first representatives. Historical resolved denotes completed computational structure prediction; two historical representatives remained unresolved. One edge-censored architecture is excluded. **C**, The two 32-member panels share six exact amino-acid sequences (18.75% of each panel; union, 58 sequences). **D**, Five previously unresolved candidates were stop-bounded on original contigs with zero extension; all five completed structural prediction and retained no clean-RT hit under unchanged criteria. The truncation conclusion applies to these five candidates. **E**, Proposed experiments test assay competence, RT catalytic dependence, array-derived cDNA and partner deletion/rescue. These experiments have not been performed.

Third, a separate *architecture-first* route inspected recurrent native locus organization within the 88,487 formal RVT-profile loci before structural prioritization. Eighteen architecture families were frozen; 17 yielded complete stop-to-stop ORFs and one edge-censored family was retained only as sensitivity. Under the same clean-RT Foldseek definition, none of the 17 complete ORF-derived representatives was strong or suggestive (one weak, 16 no-hit). Five candidates that had previously remained unresolved because of local sequence boundaries were traced back to original contigs; all five were complete stop-bounded ORFs requiring zero extension, all five yielded predicted structures, and all five remained clean-RT no-hit. For these five audited candidates, the no-hit result cannot be attributed to ORF truncation; completeness does not establish prediction accuracy or biological activity. It does not establish that the 17 proteins are a new RT family; rather, it shows that the architecture-first genomic class is not defined by recovery of a known clean-RT fold.

### Global public auditing separates sequence novelty from recognition novelty

A dated NCBI nr/nt audit found high-identity near-full public protein matches for all five final RT representatives and near-full public nucleotide matches for all five architecture regions [45]. These loci are therefore not globally novel at the sequence level. Conversely, none was compatible with the currently reconstructable exact public ART8 reference set under the frozen sequence/architecture criteria; the non-machine-readable remainder of the reported ART95 catalogue prevents an exhaustive negative crosswalk.

The result reveals a different novelty axis: *sequence deposition is not equivalent to system recognition*. A locus can be publicly deposited and even contain an annotated RT while the repeat–RT–partner relationship remains uninterpreted. MarsHill provides a historical precedent for that gap [21, 22]. Our five systems are therefore described as publicly deposited loci independently prioritized as ART-like architectures, not globally novel sequences and not proven ARTs omitted by another study (Supplementary Fig. S3).

## Discussion

Our results show that mathematical features of DNA organization can nominate genomic regions for further study without an initial protein-family label. Systematic screening applies the same prespecified scoring rules to every eligible raw-DNA window, rather than relying on retrospective recognition of unusual loci. Candidate rankings are fixed before biological interpretation, allowing both recovered and missed loci to be examined.

The benchmark provides a direct comparison with RT-directed searches. When investigators already know that RT is the relevant target class, both primary RT-plus-recurrence comparators localized all eight predeclared cases with MRR 0.875, and both secondary RVT-only comparators ranked every case first. Arm M was less effective than the RT-directed searches at ranking known RT-associated loci, but recovered all seven exact-array truths without RT-specific input. Its ranking improved when ACNV and recurrence converged. The small, ascertained benchmark supports this methodological contrast; it does not estimate performance over an unselected genome population. Arm M is therefore most relevant when the protein family or mechanism associated with an architecture is not known in advance.

The UG27 controls further test whether dual-arm convergence occurs whenever an RT is associated with an RNA array. All five UG27 systems retained exact-unit ACNV support, demonstrating that their arrays inhabit unusual positional organization, but only one of five acquired the independent recurrence support needed for dual-arm convergence. The 1/5 result was unchanged when evaluation used 28 individually mapped ncRNA units from the published annotations in place of the broader 4–5-kb experimental cassette intervals, indicating that broad locus coordinates do not explain the difference. Together with the five illustrative ART loci and 0/3 dual-arm non-array RTs, UG27 defines a descriptive specificity gradient consistent with sensitivity to a narrower recurrent DNA architecture rather than to RT identity or secondary-structure-defined RNA arrays in general.

MarsHill was a development locus, PB50 remained the predeclared Arm-M miss, and non-array RT controls showed ACNV signal without recurrence convergence. Many mathematical hits also corresponded to non-RT architectures. Retaining these results distinguishes the prespecified screening procedure from interpretations made after examining the loci.

Arm R does not require gene annotation, but its six-frame search still depends on the frozen RT profiles used to select proteins for further analysis. Structural similarity supplies fold evidence rather than enzymatic activity. For that reason, the five final architectures remain computational systems until causal assays establish RNA-dependent DNA synthesis, native array transcription, use of array RNA as substrate and partner dependence.

The independent 54.38-Gb analysis also showed that structural-compatibility fractions depend on candidate selection. Under a strict historical-rule mirror, 24/32 representatives retained at least suggestive clean-RT structural compatibility, compared with 28/30 historical resolved representatives. Applying an additional old–delta novelty filter before the same selection budget reduced this to 12/32, and the two 32-member panels shared only six exact amino-acid sequences. Thus the lower 12/32 fraction is strongly conditioned by novelty-oriented selection and does not estimate the structural-compatibility rate of the full candidate population. The matched result is substantial but attenuated transfer, while novelty-only filtering selects a different protein population.

The architecture-first route revealed a further limitation. Seventeen complete stop-to-stop ORF-derived representatives nominated by recurrent locus organization yielded no strong or suggestive clean-RT structural matches, and the five previously boundary-unresolved cases remained 5/5 no-hit after original-contig completeness was proven. These results do not establish that the 17 candidates are novel RTs or members of a single new protein family. The recurrent array–candidate–partner organizations nevertheless provide architecture-level candidates within profile-exposed loci, even without a clean-RT structural match. Biochemical experiments are needed to test whether these candidates participate in RNA-dependent DNA synthesis.

Establishing RNA-dependent DNA synthesis from array RNA, followed by partner dependence, would be necessary to connect these computationally recognized systems to a molecular mechanism. The present results support mathematical analysis of raw DNA as a complementary way to identify recurrent genomic organization before the relevant protein class is known. Proposed experimental designs for the five 6.12-Gb systems and the independent 54-Gb candidate panels are provided in Supplementary Information, Section 14, including the prospective 54-Gb extension. These experiments have not been performed; the present claims remain computational.

## Online Methods

### Raw-DNA scoring

ACNV was evaluated on both orientations of windows of 128, 256, 512, 1,024, 2,048 and 4,096 bp with stride 0.25*L*. ACNV family and recurrence were scored independently. Windows with empirical *q* ≤ 0.005 could propose regions, and strictly overlapping proposals were merged within each sequence; intervals meeting only at an endpoint remained separate. No top-*K* survival cutoff was applied.

### ACNV family

The frozen family comprises ACNV92 plus 4-, 5- and 6-mer specialist blocks. The maximum member surprisal at each window was empirically re-ranked within that scored sequence and window length. The six-length union and resulting region priority were not globally FDR-calibrated. Historical cumulative vectors are retained as sensitivity analyses rather than independent votes [34].

### Recurrence

Exact *k*-mer recurrence was scanned for *k ∈* {4, 6, 8} and lags 8–512 bp, requiring at least 32 valid pairs and two exact matches. A local-composition normalization formed the standardized statistic, which was empirically calibrated within window length.

### Formal baseline

The predeclared primary contrasts were Arm M, Prodigal–RVT plus frozen recurrence, and six-frame–RVT plus frozen recurrence. Prodigal–RVT alone and six-frame–RVT alone were predeclared secondary comparators. Prodigal used metagenomic mode followed by frozen RVT-All HMM screening [38, 39]. Overlapping same-strand HMM-positive peptides were collapsed into RT loci without a tunable gap. System localization used predeclared *±* 10-kb RT neighborhoods. Candidate scores were frozen before the benchmark truth seal was opened. The complete three-way primary results and case-level records were independently reevaluated from the frozen canonical region and RT-locus boards against sealed post-freeze truth. This evaluation reproduced the archived results without rerunning genome-wide feature extraction or HMM searches. Optional repeat-caller variants were secondary.

### UG27 reference annotations

Five experimental systems were scored genome-wide before their reference locus annotations were accessed. Individual ncRNA annotations were subsequently obtained from the published, machine-readable example-locus records and uniquely mapped into independently frozen native experimental cassettes. This annotation mapping for five systems does not reproduce the original study’s complete genome-wide covariance-model catalogue. Detector regions, scores and thresholds were not used to define the reference ncRNA intervals.

### Annotation-independent protein-space search

Raw external DNA was translated into maximal stop-to-stop segments in six frames with a minimum length of 45 aa. Frozen myRT profiles were searched with HMMER; exact-AA dereplication and family compression used MMseqs2; structural prediction and search used Boltz-2 and Foldseek; US-align was specified as an orthogonal protocol option, but available records do not establish its execution for every query; PHROG context annotation was applied only after discovery/architecture boards were frozen [41–44, 46, 47].

### Independent 54-Gb route definitions

The strict-mirror panel applies the historical four-axis selector directly to all native operational families in the independent universe and does not use a combined old–delta novelty filter. The novelty-only protein panel first excludes old–delta mixed families after joint clustering and then applies the same selector and compute budget. The architecture-first route examines the formal RVT-profile locus pool, freezes recurrent native architectures before structural ranking, and then evaluates complete stop-to-stop representatives using the same structural definition. Membership, thresholds and reference sets were not retuned after structural outcomes were observed.

### Reference-set structural compatibility

For each predicted query structure *q*, every Foldseek match *h ∈ H*(*q*) against the fixed clean-RT structural reference set was evaluated independently. Define

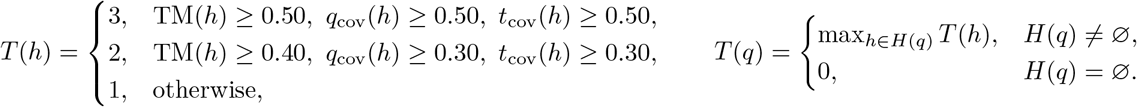

The four query-level states *T* (*q*) = 3, 2, 1, 0 are denoted strong, suggestive, weak and no-hit, respectively. The highest-TM alignment within the assigned tier is retained only as the representative alignment; it does not determine the query tier. The thresholds, clean-RT reference-set definition and Foldseek search settings were held fixed across calibration, historical, strict-mirror, novelty-only and architecture-first analyses; candidate membership within each route was frozen before structural outcomes were inspected. Independent reaggregation of the archived target-panel full-hit files reproduced the reported query tiers across 114 panel memberships, and all 112 completed query CIFs were checked against their frozen amino-acid sequences. The reaggregation did not rerun structure prediction or database searches, and the complete reference database was not independently rehashed.

### Public sequence comparison

312 The five final systems were queried against dated NCBI nr/nt. Public sequence presence was treated as deposition evidence rather than evidence that the system-level architecture had previously been recognized. Full provenance, claim rules, benchmark tables, exact UG27-unit tables, the frozen structural-compatibility definition and the completed 54-Gb delta summary are provided in Supplementary Information.

## Data availability

The data supporting the findings of this study are provided in the Supplementary Information and accompanying data tables. Public genome and protein sequences were obtained from the databases cited in the manuscript, and the corresponding accession numbers are reported in the supplementary materials. Third-party sequence and structure databases remain available from their original providers.

## Code availability

Source code associated with this study is available at https://github.com/karlieswift/ART-MathDiscovery. The repository includes analysis scripts, configuration files and tests. External genome, protein and structure databases are not redistributed and should be obtained from their original providers.

### Funding

This work was supported by the National Natural Science Foundation of China (Grant No. 12171275), the Tsinghua University Education Foundation, and the Beijing Natural Science Foundation (International Scientists Project, Grant Nos. IS25032 and IS25081).

### Author Contributions

Guoqing Hu: Conceptualization, Methodology, Investigation, Software, Formal analysis, Writing–original draft. Hao Wang: Data curation, Writing–original draft, Writing–review & editing, and submission preparation. Stephen S.-T. Yau: Conceptualization, Supervision, Writing–review & editing. All authors contributed to the manuscript and approved the final version.

