## Supplementary Information for "Deterministic DNA-first discovery reveals recurrent genomic architectures"

### 1 Evidence levels and interpretation

We distinguish five evidence levels: (1) mathematical locus nomination; (2) system architecture reconstruction; (3) protein-family/structure support; (4) biochemical activity; and (5) biological function. A locus may advance computationally without being described as a confirmed ART. In particular, high sequence identity to public nr/nt is evidence of deposition, while a repeat–RT–partner relationship is a system-level interpretation. Neither implies biochemical activity.

| Level | Evidence | Interpretation |
|---|---|---|
| L1 | ACNV/recurrence region board | mathematical candidate region |
| L2 | repeat + RT-like protein + partner candidate | computational ART-like system |
| L3 | cross-accession conservation / structure | high-priority computational system |
| L4 | RNA-dependent DNA synthesis; native substrate | biochemically validated RT system |
| L5 | partner causality + native phenotype | mechanistically characterized biological system |

### 2 Historical provenance and blinding design

The study deliberately retains the chronology of method development. The original MarsHill strict blind test used a same-window consensus and narrowly missed the predeclared Top-0.5% criterion. After unblinding, the reason was diagnosed as a multiscale mismatch: the strongest ACNV and recurrence evidence occupied different windows inside the same locus. The final region-integration procedure therefore merges proposals meeting the frozen empirical-tail threshold by coordinate overlap rather than requiring same-window coincidence. External validation genomes were not used to retune thresholds.

The frozen clean two-arm ranks are:

| Case | Role | Clean region rank | Notes |
|---|---|---|---|
| L0050 | locked retrospective | 1/84 | published 2.9-kb upstream ROI reconstructed after score freeze |
| MarsHill | development | 2/84 | development status stated explicitly |
| Madawaska | external | 2/82 | untouched external score board |
| SA1 | external | 3/78 | untouched external score board |
| Machias | external | 4/80 | untouched external score board |

### 3 ACNV mathematical formulation and order-specific family

#### 3.1 Exact frozen positional representation

The representation follows the ACNV lineage [1]; the equations below specify the frozen implementation used for this study. Its components use absolute positional products rather than a generic signed covariance formula. Let $S = s_1 \dots s_L$ be one oriented DNA window, with local positions numbered $1, \dots, L$. For $b \in \{A, C, G, T\}$, define

$$n_b = \sum_{j=1}^{L} \mathbf{1}[s_j = b], \qquad \mu_b = \begin{cases} n_b^{-1} \sum_{j:s_j=b} j, & n_b > 0, \\ 0, & n_b = 0. \end{cases} \tag{1}$$

For a token $t = t_1 \dots t_k$, let $O_t = \{i : s_i \dots s_{i+k-1} = t\}$ be its occurrence starts and $U_t = \bigcup_{i \in O_t} \{i, \dots, i+k-1\}$ the set of positions covered by at least one occurrence. The implemented component is

$$C_t(S) = \begin{cases} \dfrac{\sum_{j \in U_t} \prod_{a=1}^{k} |j - \mu_{t_a}|}{\prod_{a=1}^{k} n_{t_a}}, & O_t \neq \varnothing, \\ 0, & O_t = \varnothing. \end{cases} \tag{2}$$

The use of a set is material: positions covered by overlapping occurrences of the same token contribute once. A repeated nucleotide in $t$ repeats its count factor in the denominator. All $4^k$ tokens are retained in deterministic A,C,G,T lexical order, including zero components for absent tokens. The $k = 1$ component is a mean absolute positional deviation. The legacy extractor also computes a second-moment vector, but the primary scanner discards that vector.

The extraction buffer concatenates four counts, four means and the complete $k = 1, \dots, 6$ blocks, giving $8 + 4 + 16 + 64 + 256 + 1024 + 4096 = 5468$ features. The four scored members are:

| Member | Features | Dimension |
|---|---|---|
| $A_0$ (ACNV92) | counts, means, $k = 1, 2, 3$ blocks | 92 |
| $A_4$ | $k = 4$ block | 256 |
| $A_5$ | $k = 5$ block | 1024 |
| $A_6$ | $k = 6$ block | 4096 |

Their zero-based buffer slices are $[0, 92)$, $[92, 348)$, $[348, 1372)$ and $[1372, 5468)$. These feature-buffer indices are distinct from genomic coordinates. A genomic window is stored as a 0-based half-open interval $[s, e)$; local position $j$ maps to $s + j - 1$ on the forward strand and $e - j$ on the reverse-complement strand.

#### 3.2 Robust scores and orientation selection

The scanner processes one eligible DNA sequence per call. Calibration is performed separately for each scored sequence, window length $L$ and, where indicated, strand $\sigma$. For separately scored assembly contigs, this is contig-level calibration. Let $x_{wj}$ be feature $j$ of window $w$ for one member and orientation. Define

$$m_j = \operatorname{median}_w(x_{wj}), \qquad d_{wj} = |x_{wj} - m_j|, \tag{3}$$

$$a_j = 1.4826 \operatorname{median}_w(d_{wj}), \qquad b_j = \operatorname{mean}_w(d_{wj}), \tag{4}$$

$$s_j = \begin{cases} a_j, & a_j > 10^{-12}, \\ b_j, & a_j \leq 10^{-12}, \end{cases} \qquad J = \{j : s_j > 10^{-12}\}. \tag{5}$$

For $J \neq \varnothing$, the member score is

$$A_{m,L,\sigma}(w) = \sqrt{\frac{1}{|J|} \sum_{j \in J} \min\left(\frac{d_{wj}}{s_j}, 20\right)^2}. \tag{6}$$

If $J$ is empty, all scores in that member stratum are zero. The feature buffer and robust deviations use float32 values; the final mean of squared clipped deviations uses float64 accumulation.

Let $\overline{r}(v; v_1, \ldots, v_n)$ denote the ascending average rank, including ties. Define the percentile and empirical upper-tail transforms as

$$P(v) = \frac{\overline{r}(v)}{n}, \qquad Q(v) = \frac{n - \overline{r}(v) + \frac{1}{2}}{n} = \frac{\#\{i : v_i > v\} + \frac{1}{2}\#\{i : v_i = v\}}{n}. \tag{7}$$

The implementation additionally enforces the lower bound $1/(2n)$, which is already attained by a unique maximum among finite values. When all values in a calibration stratum are tied, every value receives $Q = 0.5$. These are empirical midrank upper-tail fractions of the observed score distribution, not FDR-adjusted values or guaranteed null-model $p$ values. The notation $q$ below refers only to this empirical fraction.

Each orientation is percentile-ranked first. With $n_L$ genomic windows of length $L$, the orientation collapse and member calibration are

$$B_{m,L}(w) = \max_{\sigma \in \{+,-\}} P_{m,L,\sigma}\big(A_{m,L,\sigma}(w)\big), \qquad q_m(w) = Q_L\big(B_{m,L}(w)\big). \tag{8}$$

Thus the selected orientation maximizes its within-orientation percentile, not the uncalibrated raw score. Average ranks resolve score ties; the first orientation is retained for equal maxima. The subscript $L$ on $Q$ means that ranking uses the genomic windows of the same length in the same scored sequence.

##### 3.3 Window-level ACNV family calibration

The maximum member surprisal is computed at each genomic window:

$$M_A(w) = \max_{m \in \{0,4,5,6\}} [-\log_{10} q_m(w)], \qquad q_A(w) = Q_L\big(M_A(w)\big). \tag{9}$$

This re-ranks the observed four-member maximum within each window length. No maximum over all six lengths is formed before calibration. The subsequent multiscale interval union and regional priority do not receive a global FDR adjustment or region-level tail recalibration. For example, the frozen tail function maps scores $(1, 1, 3, 4)$ to $(0.75, 0.75, 0.375, 0.125)$, illustrating both the average-tie rule and the finite-sample floor.

##### 3.4 Historical cumulative representations

Historical cumulative representations, including ACNV-1368, remain part of the development record and sensitivity analyses. The 1368-dimensional construction collects four counts, four means and the $k = 2, 3, 4, 5$ blocks; it is distinct from the primary extraction buffer and is not an additional proposal vote. In MarsHill development the best truth-local ranks progressed from 24/16,390 (ACNV92) to 20/16,390 (4-mer), 16/16,390 (5-mer) and 12/16,390 (6-mer). These development results motivated retaining the high-order specialist; MarsHill is not relabelled as an independent validation case.

#### 4 Generic positional recurrence

For a forward-oriented genomic window, $k \in \{4, 6, 8\}$ and integer lag $d \in [8, 512]$, define $n_{k,d} = L - k + 1 - d$ and

$$M_{k,d} = \sum_{i=1}^{n_{k,d}} \mathbf{1}[s_i \ldots s_{i+k-1} = s_{i+d} \ldots s_{i+d+k-1}]. \tag{10}$$

Only pairs with $n_{k,d} \geq 32$ are evaluated. With nucleotide frequencies $p_b = n_b/L$, the composition normalization is $p_k^{(0)} = (\sum_b p_b^2)^k$, clipped to $[10^{-12}, 1 - 10^{-12}]$. The standardized value is

$$Z_{k,d} = \begin{cases} \max\left(0, \dfrac{M_{k,d} - n_{k,d} p_k^{(0)}}{\sqrt{n_{k,d} p_k^{(0)} (1 - p_k^{(0)}) + 10^{-9}}}\right), & M_{k,d} \geq 2, \\ 0, & M_{k,d} < 2. \end{cases} \tag{11}$$

The recurrence score and empirical tail are $R(w) = \max_{k,d} Z_{k,d}$ and $q_R(w) = Q_L(R(w))$. The maximum over $k$ and lag therefore precedes within-length calibration. The binomial-style variance is a normalization; overlapping sequence pairs are dependent, so the formula does not establish a standard-normal testing distribution. No ART-specific motif or lag band is provided. Tied positive maxima retain the first pair in increasing $k$ and increasing $d$; if every score is zero, the recorded best $k$ and lag remain zero.

### 5 Integration and ranking of ACNV and recurrence regions

The frozen window lengths are $\{128, 256, 512, 1024, 2048, 4096\}$ bp, with stride $L/4$. A terminal window is appended when the regular grid does not end at the sequence boundary. The audited primary scorer rejects non-ACGT input rather than silently deleting bases. The archived primary scan also emitted an irregularity family and a historical three-family board. The final clean replay reconstructs the board solely from the frozen $q_A$ and $q_R$ columns; irregularity contributes neither proposals nor priority votes and the feature scores are not recomputed.

Each of the two families proposes windows when $q_f(w) \le \alpha = 0.005$. Within each sequence, proposals are merged into connected components under strict interval overlap. Intervals meeting only at an endpoint remain separate. All resulting union regions are retained, with no additional top-$K$ survival cutoff. The empirical-tail threshold remains an upstream selection step.

For a region $r$ and family $f$, let

$$b_f(r) = \min_{w: w \cap r \neq \varnothing} q_f(w), \qquad c(r) = \#\{f : \text{a selected proposal of } f \text{ overlaps } r\}. \tag{12}$$

The minimum uses all frozen windows overlapping $r$, including windows that did not individually pass the proposal threshold. Define

$$S(r) = -\log_{10} b_A(r) - \log_{10} b_R(r), \qquad b_{\min}(r) = \min\{b_A(r), b_R(r)\}. \tag{13}$$

The deterministic ordering is $c(r)$ descending, $S(r)$ descending, $b_{\min}(r)$ ascending and genomic start ascending. The two family minima may occur at different positions or lengths; $c(r) = 2$ defines a dual-arm region. The two views are calibrated separately, but their statistical independence is not established. $S(r)$ is a priority score, not a combined $p$ value. Neither the merged regions nor the reported priority order are recalibrated as a global error rate. The UG27 adapter enforces one scoring contig per board before assembling summary results.

### 6 Sequence-based verification of known ART loci

L0050 was localized on the complete 228,907-bp Logan contig `SRR12632059_735`. The published 2.9-kb upstream ROI was reconstructed as $[153152, 156052)$ (0-based, half-open) and lay within frozen region AR0001 $[153088, 157696)$. The recovered input FASTA and complete 14,069-window score matrix match their frozen file checksums. Independently recalculating both within-length empirical tails reproduced all 28,138 $q$ values; applying the frozen threshold, strict-overlap merge and ranking rules reproduced all 137 selected proposals and 84 union regions. Direct recurrence calculation from DNA $[155136, 155392)$ reproduced $k = 8$, lag 154 bp, 15 exact matches among 95 eligible pairs and a standardized statistic of 271.70795. This checks the reported local numerical readout without re-extracting genome-wide ACNV features.

Earlier post-score descriptions report 14 repeat copies with 16-nt majority consensus `CATGTGTATCGCATGT` and median start-to-start spacing of 154 bp. The original copy intervals, alignment and reconstruction definition are unavailable. The consensus is therefore a descriptive historical record rather than an independently reproduced copy-level annotation; absence of a literal consensus occurrence does not by itself disprove a majority consensus. The validated 154-bp recurrence maximum remains supported directly by the source DNA.

MarsHill uses the frozen upstream array zone $[213847, 215071)$ (0-based, half-open), upstream of reported gp206 RT CDS 215089–216594 and gp207 CDS 216596–217420. In the 24 October 2026 post-freeze native-DNA audit of `MW248466.1`, five nonidentical 18-nt anchors were re-extracted at start0 positions 213994, 214207, 214404, 214614 and 214787. Start-to-start spacings were 213, 197, 210 and 173 bp; their 18-nt majority consensus is `ATATGAATACGTATAATG`. These observations were derived after mathematical score freeze and did not enter locus ranking. Madawaska, Machias and SA1 strict 15-nt anchors are shown in Main Fig. 3 and Supplementary Table S2. The original CDS annotations retain their source conventions; the majority consensus is descriptive, not a universal ART motif. The native-DNA re-extraction does not establish recovery of the original historical anchor-annotation files or their full provenance.

### 7 Frozen conventional-method benchmark

#### 7.1 Design

The primary positive panel was frozen as Madawaska, Machias, SA1, LY01, S6, PALS_2, UFV_DC4 and PB50. MarsHill remained development-only and L0050 a locked retrospective sensitivity case. The dated benchmark contract defines Arm M, Prodigal–RVT plus frozen recurrence, and six-frame–RVT plus frozen recurrence as the primary contrasts. Prodigal–RVT alone and six-frame–RVT alone are predeclared secondary comparators. Prodigal used metagenomic mode and the same frozen RVT-All profile collection. HMM-positive calls were collapsed into same-strand overlapping RT loci with gap tolerance zero before truth opening. The primary neighborhood was ±10 kb; ±5 and ±20 kb were sensitivity-only. Recurrence was read from the already frozen recurrence board so that comparison isolated DNA-first versus RT-first exposure rather than confounding the comparison with a weaker repeat detector.

#### 7.2 Primary results

**Supplementary Table S3 | Frozen conventional-method benchmark.** The three primary contrasts are reported on the predeclared eight-case panel. PB50 remains a failure of Arm M; the seven-case exact-array subset is reported separately.

| Endpoint | Arm M full | Prodigal–RVT + recurrence | Six-frame–RVT + recurrence |
|---|---|---|---|
| Primary eight recovered | 7/8 | 8/8 | 8/8 |
| Hit@1 | 1/8 | 6/8 | 6/8 |
| Hit@3 | 6/8 | 8/8 | 8/8 |
| Hit@5 | 7/8 | 8/8 | 8/8 |
| MRR | 0.406 | 0.875 | 0.875 |
| Exact-array seven recovered | 7/7 | 7/7 | 7/7 |
| Exact-array Hit@3 | 6/7 | 7/7 | 7/7 |
| Exact-array MRR | 0.464 | 0.929 | 0.929 |
| Strict external array subset | 3/3 | 3/3 | 3/3 |

The corresponding predeclared secondary Prodigal–RVT-only and six-frame–RVT-only comparators each achieved eight-case Hit@1 8/8 and MRR 1.000. PB50 has RT-proxy truth rather than exact array coordinates; this determines the separately reported exact-array denominator without removing its Arm-M failure from the primary panel.

**Case-level ranks for the three primary contrasts.** A dash denotes no overlap with the truth interval anywhere on the frozen candidate list, contributing zero reciprocal rank.

| Case | Arm M full | Prodigal–RVT + recurrence | Six-frame–RVT + recurrence |
|---|---|---|---|
| Madawaska | 2 | 1 | 1 |
| Machias | 4 | 2 | 2 |
| SA1 | 3 | 1 | 1 |
| LY01 | 1 | 1 | 1 |
| S6 | 3 | 1 | 1 |
| PALS_2 | 2 | 1 | 1 |
| UFV_DC4 | 3 | 1 | 1 |
| PB50 | – | 2 | 2 |

All 77 archived case–method records were independently recalculated from the received canonical candidate boards and RT-locus tables. All 1,386 compared result fields agreed, including local recurrence ranks for 47 RT neighborhoods. The truth intervals for all eight primary cases and the two excluded development/retrospective cases matched their original source rows. The three non-array controls were evaluated against the received sealed final truth table; their separate upstream annotation table was unavailable. SD48–SD53 supply truth, case-level results, summaries, uncertainty and source-board availability. A further independent reconstruction from the archived HMMER domain tables and peptide-coordinate headers reproduced all 53 accepted RT candidates and 53 collapsed RT loci across the 13 cases; SD57–SD58 retain these checks. This is replay of archived outputs, without a new genome-wide feature scan or HMM search.

#### 7.3 Mathematical-view decomposition

On the seven exact-array cases, the full coordinate-based integration procedure recovered 7/7, with Hit@3 6/7, Hit@5 7/7 and MRR 0.464. ACNV alone and recurrence alone each touched 7/7 truths somewhere on much larger candidate lists but ranked them poorly: ACNV-only Hit@3 0/7, Hit@5 0/7, MRR 0.084; recurrence-only Hit@3 1/7, Hit@5 1/7, MRR 0.151. Thus convergence improves prioritization rather than merely unioning two independent sensitivity lists.

Adding recurrence to either target-aware RT-first search did not improve ranking. Both RVT-only routes achieved eight-case Hit@1 8/8 and MRR 1.000, whereas their recurrence-augmented counterparts achieved Hit@1 6/8 and MRR 0.875. On the exact-array seven, the corresponding MRRs were 1.000 and 0.929. This negative result supports the interpretation that recurrence is not a generic RT-HMM reranker; its utility arises when it converges with an independent mathematical architecture signal in whole-genome search.

A targeted post-freeze sensitivity analysis fixed proposal thresholds $q = 0.0025, 0.005, 0.01$ before computation; the formal threshold remained $q = 0.005$. Complete archived window matrices were available for four of the original 13 benchmark cases (L0050, Madawaska, Machias and SA1) and two previously exposed, unlabelled background inputs (NC_002666.1 and NC_028671.2). The analysis covered 18 of the 45 prespecified case–threshold combinations; the other 27 were not computed because nine benchmark cases lacked complete matrices. For the three thresholds in that order, candidate-region counts were 48/84/125, 43/82/127, 47/80/136 and 46/78/132, respectively. The corresponding best historical-target ranks were 1/1/1, 2/2/2, 6/4/6 and 1/3/5. All four targets overlapped a retained region at every threshold, but prioritization was not uniformly stable. The two background inputs yielded 6/11/24 and 6/14/25 regions and were not treated as certified negatives. At the formal threshold, all region ranks and coordinates reproduced the original canonical lists. Only the proposal threshold varied; the complete window inventory, empirical calibration, recurrence, merging and ranking functions were retained. No threshold was selected from these results. This is historical sensitivity analysis, not new external validation, and the empirical $q$ values are not global false-discovery-rate estimates.

Six separately prespecified engineering runs recomputed ACNV and recurrence scores without feature-cache reuse for native SA1 prefixes of 8,192, 32,768 and 65,536 bp, using two or four CPU workers. Two/four-worker elapsed times were 15.672/9.397, 56.867/33.967 and 124.863/69.267 s, respectively; the complete output-matrix hashes were identical between worker counts at each length. Each operating point had one trial on a shared CPU host with an eight-core quota and an 8-GiB memory limit. Timing included feature extraction and calibration but excluded later region-board construction and RT-based routes. Reported memory is maximum-child RSS, not summed concurrent-worker memory; the initial process-tree memory observer was unavailable, so aggregate RSS/PSS is missing. These limited measurements do not establish a general speedup or a 54-Gb mathematical-scan runtime.

#### 7.4 Candidate burden and exposure

On the primary eight-case panel, mean candidate burden was 301.24 mathematical union regions/Mb versus 5.13 RT loci/Mb for each protein-first route. Mean genome exposure was 100% for Arm M, 10.647% for Prodigal–RVT and 10.658% for six-frame–RVT. On the seven exact-array cases, the corresponding candidate burdens were 301.99 regions/Mb and 4.82 loci/Mb, while the mean actionable dual-arm burden was 24.75 regions/Mb (46 dual-arm regions in total). These are descriptive review workloads, not false-positive rates, because candidate units and exposure strategies differ. In bacterial controls, ±10-kb RT neighborhoods exposed only a small fraction of each genome (Prodigal: about 0.43% for Sen2, 2.93% for Ec78/Eco7 and 0.79% for DRT2), whereas Arm M exposes the entire genome mathematically.

#### 7.5 Descriptive uncertainty on the primary panel

After result freeze, exact empirical case-bootstrap percentile intervals were calculated over all $8^8$ ordered resamples using multinomial weights. The 95% intervals for eight-case MRR were 0.240–0.604 for Arm M and 0.688–1.000 for each primary RT-plus-recurrence comparator. Preserving case pairing gave an Arm-M-minus-comparator mean reciprocal-rank difference of $-0.469$, with interval $[-0.604, -0.302]$, for each comparator. These intervals describe resampling uncertainty conditional on these eight ascertained cases; they do not supply population specificity, unselected-genome accuracy or a transportability guarantee. No scores, truth coordinates or decision thresholds changed during this post-freeze calculation.

### 8 Specificity controls and public ART source panel

Sen2, Ec78/Eco7 and DRT2 are bona fide RT-containing systems but none is dual-arm positive. Their truth loci overlap mathematical union regions and can be ACNV-anomalous, but recurrence does not meet the frozen proposal threshold. Their union-board ranks were 186/1477, 350/1493 and 188/1636. This result is therefore not equivalent to "Arm M rejects RTs"; it states that RT identity alone does not create the independent recurrence signal.

Eight NCBI-resolvable public ART-related sources were retained as an exact-source crosswalk. Seven author-explicit array loci localized under the frozen mathematical engine. PB50 remained a locus-level miss despite strong mathematical regions elsewhere in its genome.

### 9 Broad 919-genome architecture discovery

The 919-genome cohort manifest matches the original frozen checksum and the declared exclusion of 61 cases from the 980-genome input set. The archived scan yielded 3,674 frozen dual-arm regions, with 3,674 unique candidate IDs from 828 genomes with at least one candidate in the recovered evidence board; 91 cohort genomes have no candidate row on that board. Every row has two-arm support, both empirical-tail values at or below 0.005, and consistent coordinates and surprisal arithmetic; all 27 mathematical fields agree with the independently archived frozen candidate table. Reapplying the archived post-score selection—zero core RT-profile loci and at least one exact or approximate repeat overlap—recovered exactly the 602 candidate IDs and strata in the original coverage table: 262 exact-repeat-linked and 340 approximate-only loci, spanning 352 genomes. These are repeat-linked architectures; absence of a core profile call does not establish absence of an RT protein.

The original Stage05E1 architecture-family program, its member and edge files, and the Stage05E2 global specificity program and outputs have now been recovered. In an isolated *post-freeze replay*, the original Stage05D1 candidate and repeat/ORF registries and Stage05D4R1 FAM50 family-compression output were supplied to the recovered E1/E2 programs, without rerunning the raw-DNA scan. E1 linked two N1/N2 candidate loci when they shared an eligible reverse-complement-canonicalized repeat-core string and a common FAM50 protein cluster whose ORF was linked within the frozen 250-bp compact radius; retained replicated connected components spanned at least two distinct source sequence IDs. Among 602 eligible candidates, 1,166 primary edges yielded 66 replicated architecture components containing 324 loci. At the prespecified 1,000-bp sensitivity radius, 1,835 edges yielded 74 replicated components containing 388 loci. E1 repeat-core string equality need not imply experimentally demonstrated or primary-exact biological repeat identity: some N2 core descriptions derive from approximate sensitivity procedures.

Stage05E2 tested signature specificity against the frozen 999-candidate Stage05D1 audit universe, not against all sequences in nature. In the strict E2A stratum, an internal E1 edge had to jointly carry a family-exclusive `EXACT_CORE_PRIMARY_DESCRIPTION` repeat core and a family-exclusive compact FAM50 cluster. Family-exclusive here is defined only within those 999 candidates, with evidence spanning at least two distinct source sequence IDs. The replay yielded 12 E2A families containing 119 member loci; the other 54 families were E2B approximate/mixed (33 families, 118 loci), E2C partially specific (20 families, 84 loci) and E2D broadly shared (one family, three loci). The archived signature table contained 179 per-candidate witness records (80 E2A and 99 E2B), so the 119 E2A family members do not correspond to 119 individually witnessed strict signature occurrences. All 15 primary E1/E2 result and gate files in the fresh upstream replay were independently confirmed to be byte-identical to the previously frozen files. These are replicated computational DNA–protein architecture components, not 66 biochemically confirmed systems or 12 nature-wide unique ART families.

SD59–SD61 retain the frozen candidate checks, 602-locus join and complete cohort mapping; the separately supplied E1/E2 replay archive retains original membership, edge, topology, witness and code/provenance records. A subsequent source-recovery audit found archived completion evidence for all 919 frozen genomes: per-genome freeze gates, selected proposals, union regions and window-score files matched 3,676 recorded expected SHA-256 values (four checks per genome). The 91 genomes with no C1 candidate rows thus have documented completed zero-dual-arm outputs, not unrun or failed scans. These are computational zero-candidate observations, not certified ART-negative genomes. Neither E1/E2 replay nor source checks independently rescan all 919 native genomes; the earlier 240-genome pilot remains provenance only.

### 10 Independent annotation-free raw-DNA protein-space discovery

#### 10.1 Raw DNA to six-frame protein universe

The initial external discovery universe contained 6.1216 Gb of raw DNA. Every sequence was translated in all six reading frames into maximal stop-to-stop segments of at least 45 aa, avoiding dependence on start-codon prediction or a gene caller. This produced approximately 65.3 million peptide segments.

#### 10.2 Frozen RT profile search and family compression

RVT screening used the 45-profile myRT collection [2] with HMMER [3]. The formal set contained 10,598 loci. Exact-AA dereplication and MMseqs2 clustering [4] produced 4,912 operational families. Thirty-two priority families were frozen across prespecified axes under a fixed structural-compute budget; the priority-panel structural positive fraction is not extrapolated to the full 4,912-family universe.

#### 10.3 Predicted structures and structural compatibility

Full-structure prediction and structure search used frozen project implementations built around Boltz-2 and Foldseek [5, 6]. A separate, prespecified post-freeze US-align comparison [7] is reported for its explicit membership scope in Section 10.4; it does not establish execution for every panel. Predicted structures were obtained for 30 of the 32 priority representatives; two remained technically unresolved. Under the fixed clean-RT reference-set definition below, 20 were strong, eight suggestive and two weak, so 28/30 representatives with completed prediction attained at least suggestive compatibility. The two unresolved representatives are not classified as structural negatives. Structure is used as fold evidence rather than as an enzymatic assay.

For each query $q$, all Foldseek matches $h \in H(q)$ against the fixed clean-RT structural reference set are evaluated independently:

$$T(h) = \begin{cases} 3, & \mathrm{TM}(h) \geq 0.50,\ q_{\mathrm{cov}}(h) \geq 0.50,\ t_{\mathrm{cov}}(h) \geq 0.50, \\ 2, & \mathrm{TM}(h) \geq 0.40,\ q_{\mathrm{cov}}(h) \geq 0.30,\ t_{\mathrm{cov}}(h) \geq 0.30, \\ 1, & \text{otherwise,} \end{cases} \qquad T(q) = \begin{cases} \max_{h \in H(q)} T(h), & H(q) \neq \varnothing, \\ 0, & H(q) = \varnothing. \end{cases}$$

Here $T(q) = 3, 2, 1, 0$ correspond to strong, suggestive, weak and no-hit. A representative alignment is selected only after tier assignment as the highest-TM match within the assigned tier. Independent reaggregation of the recovered original R5C hit tables, excluding same-accession self hits for known RTs, yielded 31 strong and one suggestive known-RT controls and no strong-or-suggestive calls among 96 matched non-RT controls (43 weak and 53 completed no-hit outcomes). The archived implementation instead graded each query by its highest-TM alignment and yielded 29 strong, one suggestive and two weak known RTs (30/32 at least suggestive); the all-hit rule changes three query tiers without changing thresholds, reference members or search settings. The earlier 32/32-strong summary was not reproduced and is corrected here. This 128-control reaggregation is separate from the 114-membership target-panel audit below. A subsequent replay used the original Foldseek protocol with the recovered source databases and independently materialized portable databases. All 128 current source amino-acid sequences were preserved exactly. The independent portable replay reproduced the complete multiset of 1,126 raw alignment rows, including 32 excluded self hits, and all control classifications; raw-file hash differences were explained by row order. Runnable 32-reference clean-RT and 96-control non-RT database subsets, component hashes and replay records are supplied. This establishes reproducibility with the recovered current inputs, without retrospectively proving the identity of historical full-database payloads lacking complete contemporaneous fingerprints.

Controls were annotation-selected and matched using Swiss-Prot sequence lengths; self-hit exclusion does not establish family-level independence. Auditing found agreement between recorded length and the recovered structural model for 95 of the 96 non-RT controls. The exception was P11578, matched to Q6A548 at a recorded length of 1,123 amino acids but represented in the recovered database by a 126-amino-acid model whose header identifies it as a fragment. The frozen 96-control panel was retained: 43 weak, 53 completed no-hit and no strong-or-suggestive outcomes. An explicitly post hoc sensitivity analysis excluding P11578 yielded 43 weak and 52 completed no-hit outcomes, with no strong-or-suggestive classification among the remaining 95 controls. Zero of 96 does not imply a zero population false-positive rate. No structural threshold, reference-set member or Foldseek search parameter is altered between the historical and independent-delta analyses.

##### 10.4 Independent audit of archived structures and full-hit classification

The audit covered 114 panel memberships: 32 historical, 32 strict mirror, 32 novelty-only, 17 complete architecture-first and one separately retained edge-censored architecture. All 32 historical memberships matched the original shortlist; its two unresolved queries were 5,120-aa and 4,803-aa proteins with recorded CUDA out-of-memory failures. All 112 completed CIF polymer sequences matched their frozen query amino-acid sequences. Reaggregation of the complete archived Foldseek outputs reproduced the historical 20 strong, eight suggestive and two weak classifications from 911 hit rows; the strict-mirror 18/6/7/1 distribution from 901 hit rows; and the novelty-only 3/9/18/2 distribution from 665 hit rows. The 17 complete architecture representatives had two total hit rows, yielding one weak and 16 no-hit queries. The separately analysed edge-censored query had 11 hit rows and remained weak. Each of the five original-contig completeness controls had an existing empty Foldseek output and a sequence-matched CIF, establishing completed no-hit outcomes rather than missing search files.

The historical aggregation correction is independently reproducible from the same 911 hits. Grading only the globally highest-TM alignment for each query reproduces the earlier 10-strong/8-suggestive/12-weak summary. Applying the stated maximum-qualifying-tier rule to all matches gives 20/8/2: six queries change from suggestive to strong, four from weak to strong and six from weak to suggestive. A highest-TM hit can fail a coverage threshold while another hit qualifies, so selecting the representative alignment before assigning the query tier loses valid evidence. The correction changes neither query membership nor numerical thresholds; the two technical failures remain outside the 30-completed denominator.

The executed strict-mirror/completeness-control wrapper and classifier match their frozen package hashes, and 37 query execution records agree with the recorded search settings and completion states. Four archived fingerprint manifests agree for six reference index/type components, and the command paths are consistent with the frozen clean-RT database definition. The historical fingerprints do not cover every database byte. In the subsequent J1 reference-database verification, current source components were hashed on the source server, the six legacy fingerprints agreed, and the materialized reference subsets were received and independently verified; the 128-control search replay is reported above. The 114-membership target-panel audit itself re-evaluates archived outputs and does not rerun prediction or reference-database search for those target queries. No additional manual structure inspection or functional validation is implied.

The prespecified post-freeze J05 comparison retained 69 memberships: 32 historical, 32 strict-mirror and five 54-Gb original-contig completeness controls. The 67 available, sequence-verified CIFs were compared against all 32 recovered clean-RT references, giving 2,144/2,144 completed US-align pairs with zero failures. The two historical CUDA-out-of-memory cases remained explicit unresolved rows. Thus all 32 strict-mirror members, including the original 18 strong and six suggestive classifications, were computationally covered. Query structures were supplied without cropping; reference C-alpha coordinates were exported by Foldseek `convert2pdb` mode 0 and checked against the current reference sequences. The frozen protocol used monomer protein alignment without a fast-mode shortcut or a TM cutoff. Inputs, executable and protocol hashes passed the final integrity check.

US-align reported separate TM scores normalized by query and reference length. For descriptive summarization, each query's best reference maximized $\min(\mathrm{TM}_q, \mathrm{TM}_r)$, breaking ties by their mean and then reference identifier. The median best minimum was 0.502010 among the 30 historical structures, 0.495765 among the 32 strict-mirror structures and 0.243830 among the five completeness controls. The complete pair matrix, original Foldseek representative-pair comparisons, aligned fractions and query-confidence summaries are supplied. Full-input comparison does not imply that every residue was aligned. These length-normalized scores differ from the original local Foldseek grading and are not a newly calibrated RT classifier; no archived tier was changed. Length and domain-coverage differences limit biological interpretation of these comparisons. No additional manual expert assessment of the 69 memberships was performed as part of this analysis. The comparison did not include new structure prediction or functional validation.

##### 10.5 Native architecture reconstruction

For structurally compatible families, native DNA context was reconstructed. Upstream repeats were called using frozen rules; downstream ORFs were enumerated without using existing functional annotation as a hard filter. Recurrent partner candidates were then assessed across family members. Five systems remained high priority; their repeat and partner features are summarized in the supplementary evidence tables below. The main manuscript summarizes the 6.12-Gb and 54-Gb routes and the proposed validation steps. PHROG annotation [8] was performed only after architecture selection.

### 11 Public sequence comparison and interpretation

Dated remote nr/nt searches showed high-identity near-full protein matches for all five final RT representatives and near-full public nucleotide matches for all five architecture regions. These loci are therefore not globally novel at the sequence level. None of the five was compatible with the exact public ART8 crosswalk under frozen criteria, but the reported ART95 remainder cannot be exhaustively checked from a complete machine-readable catalogue. These findings support describing the candidates as "publicly deposited loci independently prioritized as ART-like systems."

The deposition audit and proposed causal sequence are shown in Supplementary Fig. S3.

### 12 UG27 evaluation against published ncRNA annotations

#### 12.1 Scoring before reference annotation access

Five experimentally tested UG27 systems—*Alistipes putredinis*, *Bacteroides coprophilus*, *Bacteroides* sp. 9-1-42FAA, *Bacteroides* sp. 2-1-16 and *Blautia hansenii*—were represented by five native assemblies and 15 contigs (approximately 20.66 Mb). Whole-genome ACNV/recurrence scoring was completed and its outputs fixed before the reference locus annotations were accessed. The final candidate-region table generated before annotation access contained 6,212 union regions and 303 dual-arm regions. No detector threshold, lag range, region rule or rank was changed after unblinding.

#### 12.2 Cassette-level overlap

Construct-to-native provenance defined the experimental cassettes independently of score. Cassette-level unblinding yielded ACNV direct overlap in 5/5 systems, clean dual-arm direct overlap in 1/5 and ACNV-only direct overlap in 4/5.

#### 12.3 Published ncRNA annotations and cassette mapping

Machine-readable ncRNA annotations were obtained from the public example-locus GenBank file accompanying the original UG27 study. Thirty-nine raw ncRNA annotations were consolidated into 31 nonredundant annotated units using an annotation-only rule: overlapping annotations were collapsed and the original study's CMsearch MW-v3 intervals were preferred over overlapping manual intervals. The complete published example-locus sequences were then aligned to independently frozen experimental cassettes without reading detector output. Twenty-eight ncRNA units lay within the cassettes and mapped uniquely with an exact sequence match: 8, 6, 6, 6 and 2 units across the five systems. The three additional ncRNA units in the published *Blautia* example locus lie outside the experimental cassette. The published *Blautia* example locus differs from the experimental-cassette boundary by one cassette-end nucleotide, but the aligned 4,329-bp block is 100% identical with zero mismatch and zero gap; both in-cassette ncRNA units map uniquely and exactly.

The reference ncRNA intervals for these five systems were derived from published, machine-readable example-locus annotations. This mapping does not reproduce the original study's complete ncRNA-calling pipeline or its genome-wide covariance-model catalogue.

#### 12.4 Evaluation against mapped ncRNA units

**Supplementary Table S4 | UG27 evaluation against mapped ncRNA units.** Published, machine-readable ncRNA annotations were mapped into the frozen experimental cassettes independently of detector output. Nearest dual-arm gaps are descriptive continuous quantities; no nearby cutoff was introduced after unblinding.

| System | Exact units | Dual units | Dual fraction | ACNV-only units | Nearest dual gap |
|---|---|---|---|---|---|
| *A. putredinis* | 8 | 3 | 0.375 | 0 | 0 |
| *B. coprophilus* | 6 | 0 | 0 | 2 | 95,405 |
| *Bacteroides* sp. 9-1-42FAA | 6 | 0 | 0 | 2 | 10,968 |
| *Bacteroides* sp. 2-1-16 | 6 | 0 | 0 | 2 | 10,062 |
| *B. hansenii* | 2 | 0 | 0 | 2 | 26,086 |

The primary endpoint was strict overlap of a frozen clean dual-arm region with at least one mapped ncRNA reference interval. One of five systems was positive. In *A. putredinis*, 3/8 exact units overlapped frozen dual-arm region AR0021 (rank 21). The four negative systems each had two exact units directly overlapped by ACNV-only regions. Nearest dual-arm distance was retained as a continuous descriptor; no post-unblinding nearby threshold was defined.

The exact-unit result reproduces the cassette-level 1/5 dual-arm outcome and therefore rules out broad cassette boundaries as its explanation. Together with the five illustrative ART loci and non-array RT 0/3 dual-arm, UG27 supplies a descriptive boundary in these ascertained case sets consistent with sensitivity to sequence-recurrent architecture rather than to RT identity or structured ncRNA arrays in general.

### 13 Independent 54-Gb expansion and route comparison

The expansion was constructed as an independent universe rather than as a simple scaled replicate of the initial 6.12-Gb discovery set. The initial universe contained 6,121,620,524 bp from 131 source accessions. The independent universe contained 54,377,819,157 bp from 254 new accessions; all 131 original source accessions were excluded before source freeze. Four broad source strata and deterministic acquisition order were preserved—viral/virome, environmental metagenome, host-associated metagenome and extreme/underrepresented—with frozen target allocations of 13.5, 18.0, 13.5 and 9.0 Gb, respectively. The realized set was 54.3778 Gb because selection stopped at accession boundaries.

The scientific processing rules were unchanged. Raw DNA was six-frame translated as maximal stop-to-stop segments under genetic code 11 with no start-codon requirement and a 45-aa minimum; formal RVT support required full-sequence $E \leq 10^{-4}$ and best-domain independent E-value $i\text{-}E \leq 10^{-4}$; exact-AA deduplication and operational-family compression used the same frozen MMseqs2 settings. The independent scan produced 88,487 formal RT-like instances, 85,503 exact-AA-unique sequences and 26,396 native operational families.

| Route | $n$ | Strong | Suggestive | Positive | Interpretation |
|---|---|---|---|---|---|
| Historical 6.12-Gb resolved | 30 | 20 | 8 | 28 (93.3%) | historical priority reference |
| 54-Gb strict mirror | 32 | 18 | 6 | 24 (75.0%) | historical family-universe definition; no combined novelty filter |
| 54-Gb novelty-only protein route | 32 | 3 | 9 | 12 (37.5%) | old–delta mixed families excluded before selection |
| 54-Gb architecture-first complete | 17 | 0 | 0 | 0 | recurrent native architecture; 1 weak and 16 no-hit |

#### 13.1 Strict-mirror selection under the historical rules

The strict mirror was constructed directly from all 26,396 native independent-universe operational families. It replayed the exact historical four-axis compute-budget rule—replication, maximum amino-acid length, formal model coverage and motif-topology, followed by deterministic backfill—without known annotation, structural outcome, retuning or the combined old–delta novelty filter. The resulting 32 representatives comprised 18 strong, six suggestive, seven weak and one no-hit under the frozen clean-RT structural definition, giving 24/32 at least suggestive compatibility. This is the appropriate historical-rule comparison to 28/30 among the 30 resolved historical priority representatives: substantial but attenuated structural transfer rather than collapse.

#### 13.2 Novelty-only protein selection

Joint old–delta clustering identified 24,810 delta-only combined families after exclusion of mixed historical–delta families. Applying the same frozen four-axis selector yielded 32 novelty-only representatives: three strong, nine suggestive, 18 weak and two no-hit, or 12/32 at least suggestive. The strict-mirror and novelty-only panels shared only six exact amino-acid sequences out of 32. Thus the novelty filter substantially changes the proteins sent to structural evaluation. The 12/32 fraction is conditional on novelty-only selection and does not estimate the structural-compatibility rate of the full 54-Gb candidate population.

##### 13.3 Architecture-first selection and structural outcomes

A separate native-architecture search inspected the 88,487 formal RVT-profile loci and froze 18 delta-only recurrent architecture families before structural prioritization. It was independent of structural outcome but retained upstream profile conditioning. Seventeen families yielded complete stop-to-stop ORFs; one edge-censored family was retained only as sensitivity. Under the identical frozen clean-RT Foldseek definition, the 17 complete ORF-derived representatives yielded zero strong, zero suggestive, one weak and 16 no-hit representatives. This does not establish 17 new RTs or one new protein family. Instead, it defines an architecture-level candidate class whose biological relevance remains experimentally unvalidated.

##### 13.4 Original-contig completeness control

Five architecture-first candidates had previously remained unresolved because translated segments might have been clipped by contig boundaries. Original-contig review confirmed that all five are complete stop-bounded ORFs with zero amino-acid extension. All five subsequently completed frozen Boltz structural prediction and returned five of five no-hit outcomes against frozen clean-RT Foldseek references. The absence of an RT-fold match in this subset is therefore not explained by ORF truncation or incomplete structural prediction.

The completeness control is a limitation/censoring test, not functional or biochemical validation. It does not demonstrate that the candidates lack biological function or establish a novel RT family.

##### 13.5 Comparison of the three 54-Gb analysis routes

| Route | Calibrated structural outcomes | Interpretation |
|---|---|---|
| Strict-mirror protein route | 18 strong + 6 suggestive / 32 | Protein-space compatibility, not 24 validated ARTs |
| Novelty-only protein route | 3 strong + 9 suggestive / 32 | Different sampling universe, not additive to strict mirror |
| Architecture-first complete ORFs | 0/17 RT-compatible | An independent architecture discovery branch; 5/5 initially unresolved are complete and no-hit |

The two 32-member panels share six exact amino-acid sequences. Structural compatibility counts are not biological systems. The independent source cohorts, selection rules, reference structures and fold thresholds remain frozen; the route-specific outcomes are reported separately and do not constitute experimental evidence.

All results in this section are computational. The lack of a calibrated RT-fold match in the architecture-first branch does not preclude other types of repeat-associated biology.

#### 14 Requirements for experimental validation

The present study reports computational prioritization of genomic architecture candidates but does not establish RT catalytic activity, native array-to-cDNA conversion, partner dependence or biological phenotype. The experiments below define the evidence needed for functional validation.

#### 14.1 Experimental evidence required for functional validation

| Tier | Key question | Essential decision criterion |
|---|---|---|
| 0. Positive-control platform | Can a published RNA-to-DNA system be reproduced locally? | UG27 positive controls before candidate negative results are interpreted |
| 1. RT chemistry | Does candidate protein catalyze RNA-dependent DNA synthesis? | WT versus catalytic-dead; template dependence and sequence-confirmed product |
| 2. Native repeat RNA | Is the native repeat/spacer array transcribed? | Independent array-RNA detection, including appropriate negative controls |
| 3. Substrate causality | Is detected DNA derived from cognate array RNA? | Sequence-resolved array RNA to RT-dependent cDNA connection |
| 4. Partner causality | Is the candidate partner necessary? | Perturbation with matched rescue where construct design permits |
| 5. Native biological function | What phenotype does the molecular system mediate? | Reproducible phenotypic tests only after mechanistic support |

#### 14.2 Candidate prioritization

For future biochemical and substrate testing, Systems 1–3 are the primary computational priorities, with Systems 4–5 as additional candidates. These priorities do not alter the frozen computational ranks or establish ART activity. The architecture-first 0/17 no-hit candidates form a separate exploratory group for which RT-specific assays may not be appropriate.

These proposed experiments are not included in the evidence reported here.

### 15 Supplementary evidence tables

The tables below summarize ACNV representations, known ART locus anatomy, the five computational candidate systems, public sequence matches and source-cohort composition. Supplementary Tables S3–S8 provide the benchmark, exact-unit, sequence-coordinate, data-index and source-evidence summaries. Supplementary Tables S1 and S2 contain the UG27 evaluation and Stage03B anchor catalogue, respectively (Section 21.2).

#### 15.1 ACNV representations and development sensitivity

| Representation | Dimension | Role in final method | MarsHill local rank |
|---|---:|---|---|
| ACNV92 | 92 | low-order primary specialist | 24 / 16,390 |
| 4-mer ACNV block | 256 | primary specialist | 20 / 16,390 |
| 5-mer ACNV block | 1,024 | primary specialist | 16 / 16,390 |
| 6-mer ACNV block | 4,096 | primary specialist | 12 / 16,390 |
| ACNV-344 (2–4mer cumulative) | 344 | sensitivity / method lineage | 20 / 16,390 |
| ACNV-1368 (2–5mer cumulative) | 1,368 | sensitivity / method lineage | 16 / 16,390 |
| ACNV-5464 (2–6mer cumulative) | 5,464 | sensitivity / method lineage | 16 / 16,390 |

The historical high-order sensitivity analyses informed method development. The final frozen ACNV family contains the four order-specific specialists A0, A4, A5 and A6 with empirical within-length calibration and a family-level maximum-null correction. Cumulative vectors are method-lineage analyses and are not counted as independent proposal votes.

#### 15.2 Known ART locus anatomy used only after score freeze

| Case | Frozen/source region | Post-freeze repeat description | Clean two-arm result |
|---|---|---|---|
| L0050 | published 2.9-kb ROI $[153152, 156052)$ | reported 14 copies; 16-nt majority consensus CATGTGTATCGCATGT; median spacing 154 bp | 1/84; recurrence lag 154 bp |
| MarsHill | array zone $[213847, 215071)$ | five nonidentical 18-nt anchors; 18-nt majority consensus ATATGAATACGTATAATG; spacings 213/197/210/173 bp; gp206 RT and gp207 partner downstream | 2/84 |
| Madawaska | strict core 213194–214001 | five strict 15-nt anchors; spacings 213/196/212/171 bp | 2/82 |
| Machias | strict core 218595–219017 | three strict anchors; spacings 211/196 bp | 4/80 |
| SA1 | strict core 9596–10371 | five strict anchors; spacings 212/171/201/176 bp | 3/78 |

#### 15.3 Five final computational candidate systems

| System | Family structural support | Family repeat/replication summary | Downstream partner summary |
|---|---|---|---|
| System 1 | STRONG; TM 0.7726; qcov 0.770; tcov 0.546 | ATGGAGTA; 3 copies; 157/157-bp spacers (165/165-bp start intervals); 3 independent near-context clusters | 125 aa; same strand; 13-bp gap; 99.2% cross-accession identity |
| System 4 | STRONG; TM 0.6832; qcov 0.678; tcov 0.834 | 12 strict loci; dominant CGGCAAGG | 419 aa; same strand; 20-bp gap; approximately 94.8% identity |
| System 2 | STRONG; TM 0.6531; qcov 0.702; tcov 0.818 | CGCCAAGA; median spacer 198.5 bp | 121 aa; 50-bp gap; 98.3% identity |
| System 5 | STRONG; TM 0.6337; qcov 0.749; tcov 0.942 | AGACTGCAT; median spacer 168.5 bp | 32-aa microprotein; 11-bp gap |
| System 3 | STRONG | ATTGAAGAA/TTGAAGAA family across multiple accessions | 126 aa; 16-bp gap; approximately 80.3% identity |

Family summaries can pool different member loci. Their dominant motif, median spacer and strongest structural alignment need not describe the same selected representative. The exact native DNA, selected RT and partner sequences, architecture-local intervals and repeat occurrences are given separately in Supplementary Data. In particular, a spacer excludes the repeat itself; for the 8-nt System 1 motif, a 157-bp spacer corresponds to a 165-bp start-to-start interval.

#### 15.4 Dated NCBI public audit

**Protein queries against nr.** Identity and query coverage are fractions.

| System | Top nr accession | Identity | Coverage | Public description |
|---|---|---|---|---|
| System 1 | WP_008642534.1 | 1.0000 | 0.9777 | reverse transcriptase/maturase family protein [Bacteroidales] |
| System 4 | DAQ13326.1 | 0.9856 | 0.9697 | MAG TPA hypothetical protein [Caudoviricetes sp.] |
| System 2 | WP_208319807.1 | 0.9952 | 1.0000 | RNA-directed DNA polymerase [Intestinimonas butyriciproducens] |
| System 5 | AFB75804.1 | 0.9907 | 0.9908 | hypothetical protein [Bacteriophage sp.] |
| System 3 | WP_234075906.1 | 0.9944 | 0.9834 | RNA-directed DNA polymerase [Bacteroidales] |

**Native architecture DNA queries against nt.**

| System | Top nt accession | Identity | Coverage |
|---|---|---|---|
| System 1 | CP103238.1 | 0.9977 | 1.0000 |
| System 4 | OP075715.1 | 0.9859 | 0.9862 |
| System 2 | BK048116.1 | 0.9878 | 1.0000 |
| System 5 | BK025545.1 | 0.9884 | 1.0000 |
| System 3 | BK053900.1 | 0.9981 | 1.0000 |

All five have high-identity public sequence/locus matches. Public deposition does not establish whether the repeat–RT–partner system was previously recognized. Source search outputs and query-level provenance are retained in SD35.

##### 15.5 Frozen 54-Gb source strata

| Stratum | Target bp | Deterministic eligible candidates | Interpretation |
|---|---|---|---|
| A | 13,500,000,000 | 7,684,536 | viral / virome |
| B | 18,000,000,000 | 57,598,045 | environmental metagenome |
| C | 13,500,000,000 | 3,409,961 | host-associated metagenome |
| D | 9,000,000,000 | 18,881 | extreme / underrepresented |

Candidate counts refer to the ordered eligible pool rather than the number of downloaded objects. Within each stratum, acquisition ended when the frozen realized-nucleotide quota was met. Deterministic selection was defined by within-stratum ordering independently of cross-stratum execution order.

#### 16 Sequence and coordinate data

The accompanying Supplementary Data contains 62 TSV files (SD00–SD61), including a catalog and data dictionary, together with FASTA files and an Excel companion. A row identifies its biological and coordinate unit: genomic locus, oriented native architecture core, translated peptide, repeat occurrence, ncRNA unit, region or operational family. These units are not interchangeable. Full sequence strings remain machine-readable instead of being compressed into a printed table.

##### 16.1 Five selected native systems

Supplementary Table S5 gives the exact coding intervals of the five selected native architectures. SD21 provides the complete DNA strings; SD03 provides the ten wild-type RT/partner protein sequences; SD22 supplies 31 individual features (ten coding features and 21 strict-repeat occurrences). Every coding interval was checked by exact translation, every repeat occurrence was checked against the supplied DNA and terminal stops were checked separately. Both a coding-end gap and a gap after the terminal stop are retained. This resolves an apparent 3-bp discrepancy without changing sequence or annotation.

Supplementary Table S5: Exact native architecture sequences and translated coding intervals for the five computational systems.

| System | Native strand | DNA (bp) | RT coding interval | Partner coding interval | RT (aa) | Partner (aa) | Gap (bp) |
|---|---|---|---|---|---|---|---|
| 1 | + | 5,648 | $[4042, 5254)$ | $[5270, 5645)$ | 404 | 125 | 13 |
| 2 | − | 4,018 | $[2351, 3599)$ | $[3652, 4015)$ | 416 | 121 | 50 |
| 3 | − | 2,136 | $[653, 1736)$ | $[1755, 2133)$ | 361 | 126 | 16 |
| 4 | + | 5,141 | $[2571, 3858)$ | $[3881, 5138)$ | 429 | 419 | 20 |
| 5 | − | 5,694 | $[4279, 5581)$ | $[5595, 5691)$ | 434 | 32 | 11 |

All intervals are 0-based and half-open on the supplied, RT-oriented native architecture FASTA records `SYSTEM_1_ARCHITECTURE_DNA` through `SYSTEM_5_ARCHITECTURE_DNA`. Both coding features run in the positive direction on these supplied records; "native strand" gives the RT orientation on the original assembly contig. Coding intervals exclude the terminal stop codon: for a coding end $e$, the stop occupies $[e, e+3)$ and the stop-inclusive ORF ends at $e+3$. Gap is measured from the end of the RT terminal stop to the partner coding start; for System 1, this is $5270 - (5254 + 3) = 13$ bp. Every coding interval translates exactly to the frozen WT peptide, and both terminal stop codons were checked. Original-contig core origins and the corresponding absolute coordinates for all 31 coding/repeat features were joined from the frozen source mappings and are supplied in SD42. This coordinate join did not re-extract the five cores from full parent-contig DNA. Full DNA and amino-acid sequences, source accessions, internal-to-public identifier crosswalks, per-sequence hashes, and the separate exact-repeat coordinates are in Supplementary Data SD02, SD03, SD21–SD22 and SD42. The original five-record architecture FASTA has SHA-256 `c496dca26651abd3f83b74ece325cb947d315a5fc11c2c880f932a1cfad75cbe`.

Supplementary Table S6: Original-contig coordinate mappings for the five selected systems.

| System | Original contig | Strand | Architecture core | RT coding | Partner coding |
|---|---|---|---|---|---|
| 1 | SRR5056912_2957 | + | $[16799, 22447)$ | $[20841, 22053)$ | $[22069, 22444)$ |
| 2 | SRR22135747_4611 | − | $[5720, 9738)$ | $[6139, 7387)$ | $[5723, 6086)$ |
| 3 | SRR32969331_101548 | − | $[2438, 4574)$ | $[2838, 3921)$ | $[2441, 2819)$ |
| 4 | ERR14033733_31035 | + | $[9879, 15020)$ | $[12450, 13737)$ | $[13760, 15017)$ |
| 5 | ERR14039848_24814 | − | $[355, 6049)$ | $[468, 1770)$ | $[358, 454)$ |

Coordinates are 0-based and half-open on the original assembly contig. RT and partner coding intervals exclude terminal stops. Negative-strand intervals remain in ascending original-contig coordinates; the supplied architecture FASTA is reverse-complemented relative to that interval. These mappings join frozen native-locus placement and namespace records; full parent-contig DNA was not re-extracted in this audit. SD42 provides source accessions, original headers and all repeat-occurrence mappings, and SD22 provides each corresponding local feature.

Supplementary Table S6 gives the corresponding original-contig placements.

Family-level summaries elsewhere in the Supplement can pool multiple native loci. For the selected System 2 DNA, the exact repeats have 211/205-bp spacers, whereas the family median is 198.5 bp. The selected System 4 DNA contains two strict arrays with motifs `AAGACCACC` and `CGGCAAGG`. The selected System 5 DNA contains `TACGGTGA`; the family-level dominant motif `AGACTGCAT` describes other members. These records remain separate in SD02, SD22 and the family evidence tables. The selected System 1 8-nt motif has 157/157-bp spacers and 165/165-bp start intervals.

#### 16.2 Candidate populations, reference annotations and source coverage

The release includes all 88,487 independent formal candidate characterization rows, all 26,396 family-priority rows, their architecture/context mappings, the strict-mirror 32 and novelty-only 32 candidate tables, all 18 architecture-selected families and the 17 complete structure representatives. The discovery universe is represented by all 10,598 locus and 4,912 family architecture summaries. Native-context FASTA is also available for 517 peptide/context records. Received original structure/search files and other source records are provided in a separately supplied raw-source archive; both this archive and the code-and-data archive are needed to replay the raw-source audits.

SD38 supplies exact amino-acid sequences for all 81 structural-panel memberships (32 strict mirror, 32 novelty-only and 17 complete architecture-first), comprising 75 unique peptide hashes. SD40–SD41 provide their coding DNA and 75 full native contigs from 57 source accessions. Independently extracting the stated 0-based, half-open intervals, reverse-complementing 35 negative-strand records and translating all 81 sequences reproduced every frozen protein; 46 memberships are on the positive strand. The full compressed source shards were not received or independently rehashed. SD39 links these panel memberships and the five selected native systems to their sequence, coordinate and evidence records. SD42 adds the five systems' original-contig core origins and absolute mappings for all 31 local coding/repeat features; this provenance join did not re-extract the five cores from complete parent-contig DNA.

All 13 frozen case-genome FASTA files are supplied with file sizes and SHA-256 checksums matching the frozen records: the eight primary ART cases, L0050, MarsHill and three non-array RT controls. The formal benchmark includes all 13 sealed reference-annotation records and 77 case–method evaluation rows, including the previously unavailable six-frame primary result. The canonical-board replay is complete for these received evaluation records; availability of earlier upstream source boards is listed separately. UG27 includes all 28 exact ncRNA-unit DNA/coordinate records, each checked against its native interval and the corresponding published annotation, plus five system summaries and all 6,212 frozen regions, of which 303 are dual-arm. SD44 identifies all 254 selected independent source accessions and their recorded compressed-file hashes, linked to the 118,940-event acquisition ledger; selected nucleotide counts sum to 54,377,819,157 bp.

The complete archived structure outputs support independent query-level grading across 114 panel memberships, including all 32 historical selection records and the two technical failures. SD45–SD47 preserve the individual query checks, 2,490 raw-hit classifications and historical tier transitions. These records link the reported summary counts to individual archived query and search outputs. It does not establish the accuracy of predicted folds, independent manual review or enzymatic activity.

SD19 retains the original source-coverage snapshot; subsequent recoveries and remaining limitations are summarized in the evidence table below. Large production inputs, some earlier upstream mathematical boards, the three controls' separate annotation table and the original L0050/MarsHill copy/anchor definitions remain outside the verified replay scope. Runnable current reference subsets are now supplied, while historical full-database identity remains unproven. Missing records are not assigned zero values or interpreted as biological negatives.

The supplementary data catalogue comprises SD00–SD61. Additional post-freeze computation archives document J1 `CONTROL_128_SEQUENCE_AND_LENGTH_AUDIT.tsv` and `SEARCH_REPLAY_REPORT.json`; J04 `SENSITIVITY_ALL_CASES.tsv` and `RESOURCE_MEASUREMENTS.tsv`; and J05 `PAIR_RESULTS.tsv`, `QUERY_SUMMARY.tsv` and `EXPERT_REVIEW_WORKSHEET.tsv`. These analysis-specific archives are separate from SD00–SD61; their run manifests, protocol hashes and completion records define the executed analyses. The J05 worksheet lists the 69 panel memberships; additional manual expert assessment was not part of the reported analysis.

Supplementary Table S7 indexes the accompanying machine-readable data.

Supplementary Table S7: Index of accompanying machine-readable Supplementary Data.

| Data ID | Rows | Contents |
|---|---:|---|
| SD01 | 13 | Known ART cases and non-array RT controls |
| SD02 | 5 | Five computational array-associated RT-like systems |
| SD03 | 10 | Exact wild-type RT and partner amino-acid sequences |
| SD04 | 32 | Strict mirror candidates and structural results |
| SD05 | 17 | Architecture-first complete ORF representatives |
| SD06 | 28 | UG27 exact ncRNA unit coordinates and overlap results |
| SD07 | 5 | UG27 array and native cassette summary |
| SD08 | 6,212 | UG27 complete pre-unblinding region board |
| SD09 | 4 | Observed structural compatibility by route |
| SD10 | 32 | Exact amino-acid panel membership overlap |
| SD11 | 8 | Formal conventional benchmark results |
| SD12 | 5 | Mathematical convergence ablation results |
| SD13 | 88,487 | Complete independent formal protein candidate characterization |
| SD14 | 26,396 | Complete independent operational-family priority table |
| SD15 | 13 | Discovery-universe analysis counts |
| SD16 | 4 | Independent-universe frozen source strata |
| SD17 | 8 | L0050 exact worked-example readout |
| SD18 | 5 | Broad919-genome branch summary |
| SD19 | 38 | Data coverage and source assets still required |
| SD20 | 37 | Data dictionary |
| SD21 | 5 | Five exact native architecture DNA sequences |
| SD22 | 31 | Five-system RT,partner and exact-repeat coordinates |
| SD23 | 517 | Independent positive-family native contexts and proteins |
| SD24 | 32 | Novelty-only32 candidate coordinates and structural tiers |
| SD25 | 18 | All18 architecture-first families and evaluation membership |
| SD26 | 28,328 | All independent repeat-array detections |
| SD27 | 12,241 | Independent nearest downstream ORFs and clusters |

| Data ID | Rows | Contents (continued) |
|---|---|---|
| SD28 | 88,487 | Independent full native-context evaluability |
| SD29 | 88,487 | Independent native and combined-family mapping |
| SD30 | 2,085 | All discovery-universe repeat arrays |
| SD31 | 10,598 | Complete10598 discovery locus architecture records |
| SD32 | 4,912 | Complete4912 discovery family architecture summary |
| SD33 | 16 | Sixteen core recurrent strict loci |
| SD34 | 7 | Seven-family computational evidence board |
| SD35 | 5 | Dated global public sequence audit |
| SD36 | 40 | Public ART8 exact-reference crosswalk |
| SD37 | 9 | Frozen formal benchmark registry |
| SD38 | 81 | Exact amino-acid sequences for all three frozen panels |
| SD39 | 86 | System and structural-panel sequence/coordinate crosswalk |
| SD40 | 81 | Structural-panel native ORF DNA and coordinates |
| SD41 | 75 | Structural-panel original native contigs |
| SD42 | 5 | Five-system original-contig coordinate mapping |
| SD43 | 28 | UG27 exact native-unit DNA |
| SD44 | 254 | Independent-universe selected source acquisition records |
| SD45 | 114 | Structural-query identity and all-hit tier audit |
| SD46 | 2,490 | All returned structural-search hits reclassified |
| SD47 | 30 | Historical tier transition audit |
| SD48 | 13 | Formal post-freeze truth and source replay |
| SD49 | 77 | Formal case-by-method benchmark replay |
| SD50 | 28 | Formal benchmark method/subset summaries |
| SD51 | 5 | Primary-case empirical bootstrap uncertainty |
| SD52 | 47 | RT-locus recurrence-neighborhood replay |
| SD53 | 22 | Earlier source-board availability |
| SD54 | 1 | L0050 verified score and local DNA readout |
| SD55 | 137 | L0050 complete rebuilt proposal board |
| SD56 | 84 | L0050 complete rebuilt union-region board |
| SD57 | 53 | Raw-HMM accepted candidate reconstruction |
| SD58 | 53 | Raw-HMM RT locus reconstruction |
| SD59 | 3,674 | Broad branch: all3674 candidate records |
| SD60 | 602 | Broad branch:602 repeat-linked candidate joins |
| SD61 | 919 | Broad branch: complete919-genome frozen cohort |

### 17 Data and code availability

The accompanying archive provides scientific source tables, frozen code, configuration files and region boards fixed before truth opening. It also includes the complete received primary-benchmark outputs, public ART mappings, exact UG27 sequences, candidate and family tables, structure and architecture records, dated public-audit outputs, source provenance, an environment specification, validation tools and a minimal worked example. Exact native five-system sequences and all 81 structural-panel membership amino-acid/coding-DNA records are included. Independent audits cover frozen-result reevaluation, source/sequence identity and archived full-hit structural grading. The separately supplied raw-source archive contains the received original structure/search and other source files; replay of those audits uses both archives. A separate read-only Stage05E1/E2 archive contains recovered original family/edge/witness records, the upstream-input replay audit, and the fixed per-family source crosswalk. SD00 indexes the machine-readable tables and SD19 records the scope of source coverage.

Large public sequence inputs remain at their original providers. The public code repository is available at `https://github.com/karlieswift/ART-MathDiscovery`; no archived DOI is specified for the accompanying source-data archives. The manuscript and finished figure files are supplied separately from the scientific code and data; figure input measurements remain in the relevant source tables.

### 18 Computational verification and limitations

The reproducibility record includes scorer versions, frozen thresholds and window scales, source/accession manifests, pre-unblinding score boards, benchmark truth-resolution records, UG27 exact-unit provenance, structure-reference and query-level grading definitions, software versions and file checksums. The provenance records distinguish checks of sequences and existing results from complete production reruns and analyses not performed. All three predeclared primary benchmark results have been independently reproduced from the received canonical outputs. Of 22 earlier upstream source boards, three original files were received and three additional boards were reconstructed byte-identically from archived matrices using their frozen expected hashes. Nineteen original files were unavailable for direct inspection, including the three represented by byte-identical reconstructions; 16 had neither original nor reconstructed files available. The three reconstructions reproduce downstream boards and do not repeat DNA feature extraction. The three controls' original annotation table remains unavailable. Six previously unavailable historical window matrices were obtained on 23 October 2026 (MarsHill, LY01, S6, PALS_2, UFV_DC4 and PB50). Across their 98,241 scored windows, an independent check verified every expected 128–4,096-bp window start at the frozen quarter-window stride, including each terminal flush window, with no duplicate coordinate key or length mismatch. The matrix coordinates are complete, but their historical source identity remains unverified because contemporaneous original matrix-content hashes were not supplied. The additional cases' $q$-sensitivity analyses were not recomputed. Three larger control matrices were located on the server but were not part of the transferred data used for these checks.

The archived target-structure audit recomputes query tiers and checks CIF sequences; it does not rerun target prediction or target-reference search. The separate 128-control searches have now been replayed against recovered and portable reference databases. Complete compressed production-source shards remain required for a full independent production rerun, and historical full-database identity is not retrospectively established. Original L0050 repeat-copy annotations remain unavailable. The five MarsHill 18-nt anchors and their spacings were verified from native DNA in the post-freeze N2 audit, although the original historical author-annotation files were unavailable for inspection. The broad-branch E1/E2 family tables, frozen rules, and complete D1/D4R1 upstream replay have now been recovered; all 15 core outputs match the original frozen files byte-for-byte. Family exclusivity remains bounded to the Stage05D1 999-candidate audit universe. A source audit located 498 out of 498 matrix-reference paths with documented historical expected hashes and independently matched those on the server; two additional large matrix files had no corresponding expected hashes. The files themselves were not all transferred or independently rerun here. The post-freeze N2 audit verified repeat-group, copy and spacer coordinates against native DNA for all 602 repeat-linked candidates (16,527 groups, 64,988 copies and 48,461 spacers), using 179 archived contexts and 423 frozen-FASTA candidates. This nucleotide and coordinate verification is separate from a fresh 919-genome scan or biochemical validation. The prespecified J05 scope is reported in Section 10.4; it does not establish universal US-align coverage or completed manual structural review. The study did not include wet-laboratory validation.

**Exploratory structured-RNA transfer evaluation (J2; separate from the ART main benchmark).** A complete 6,722,539-bp NC_012660.1 native genome was scored through the frozen two-arm mathematics and the four target-aware RVT comparators. Evaluation used 40 author-defined RF01731 ncRNA intervals on that one genome; computational score files were locked before the truth was opened to the evaluator, and all five method attempts completed. Strict interval overlap by Arm M recovered 33/40 sites (82.5%), with Hit@1 = 0/40, Hit@3 = 2/40, Hit@5 = 3/40 and mean reciprocal rank = 0.04624 over 2,025 whole-genome candidate regions (301.23 candidates/Mb). Each RVT route produced one neighborhood proposal and overlapped 0/40 structured-RNA intervals. Because these author-defined RNAs are not required to encode or neighbor RTs, this is not evidence of ART-method superiority or an empirical false-positive-rate comparison. All 40 intervals belong to one genome/one independence unit; no independent ART-primary evaluation, cluster-bootstrap interval or population specificity estimate is claimed. The author source belongs to a literature family already encountered during UG27 analysis, and source-exposure and family-disjoint generalization remain uncertain. The annotation crosswalk and biological interpretation have not been independently validated. The exact post-freeze per-target ranks, candidate burdens and independently reaggregated summaries are retained as separate J2 audit files without modifying SD00–SD61 or the frozen primary benchmark.

Supplementary Table S8 summarizes the computational verification and source-data limitations for the ten evidence items tracked in SD19, including the distinction between computational checks and experimental validation.

Supplementary Table S8: Computational verification and source-data limitations for the ten evidence items tracked in SD19.

| ID | Source evidence and verification | Relevant manuscript sections | Reproducibility and interpretation limits |
|---|---|---|---|
| P01 | Archived scan-completion verification: all 919 genomes have verified C1 completion plus four recorded input/output hashes each (3,676/3,676 checks). Exactly 91 completed genomes produced zero C1 dual-arm candidate regions; these are not biological negatives. | Main Fig. 4; S9 and S16. | Archive-file identities are verified; a fresh independent 919-genome rerun was not performed. |
| P02 | Frozen E1 computational verification: original 66-family memberships, 1,166 edges and rules recovered; upstream D1/D4R1 replay reproduced 66 families/324 loci, with original result files byte-identical. | Main Fig. 4; S9. | Frozen E1 replay is complete, and N2 subsequently verified native repeat-coordinate provenance; biological interpretation remains distinct from biochemical validation. |
| P03 | Frozen E2 computational verification: original 12 E2A-family memberships (119 loci), exact-primary/FAM50 criteria, and 179 witness records recovered; replayed E2 outputs are byte-identical. | Main Fig. 4; S9. | Family-exclusive means exclusive in the frozen 999-candidate universe, not natural uniqueness or ART function; the 119 family loci include 80 directly witnessed E2A candidate records. |
| P04 | Original server retains 498/498 matrix paths with known expected hashes; two additional large files lack expected hashes. Six distinct missing historical case matrices were transferred and their 98,241 quarter-stride window keys validated, but no matching original frozen matrix-content hashes were available. | Main Online Methods; S3–S7 and S16. | Historical source identity and original computational settings are not fully verified for all matrices. The additional $q$-threshold sensitivity cases were not recomputed; file presence alone does not establish a full production replay. |
| P05 | Control truth provenance: sealed final intervals are available for all three controls; their single shared upstream annotation table is absent. | Main Table 1; S7–S8. | SD19 records the frozen SHA-256 for `C1BO_CONTROL_TRUTH_RESOLVED.tsv`. The separate upstream annotation provenance cannot be fully reconstructed without the missing annotation table. |
| P06 | Earlier mathematical source boards: 3/22 original files received; three more boards reconstructed byte-identically from archived matrices; 16 lack either form. Canonical B0 boards remain complete. | Main Table 1; S7 and S16. | Nineteen original files were unavailable for direct inspection; three have exact-hash reconstructions, while 16 have neither original files nor reconstructions available. |
| P07 | L0050 repeat-copy reconstruction: original 14-copy intervals/alignment absent; the score matrix, region replay and 154-bp recurrence result are verified. | Main Figs. 2–3; S6 and S15. | The original per-copy intervals, alignment and consensus definition for the reported 14-copy reconstruction were unavailable. |
| P08 | N2 verified five MarsHill 18-nt anchors, their coordinates and spacings on frozen native DNA. | Main Fig. 3; S6 and S15. | Native-DNA verification is complete; the original historical author-annotation files were unavailable for inspection. |
| P09 | Runnable clean-RT (32-reference) and non-RT (96-control) database subsets, hashes and exact replay records are supplied. Current sequence identity is verified; one model-length exception is disclosed. | Main Fig. 5; Supplementary Fig. S3; S10 and S13. | Historical database identity is not established without contemporaneous full-payload fingerprints or an authenticated historical copy. Replay is supported for the current inputs. |
| P10 | The archived N2 audit (24 October 2026) reports completed native-DNA coordinate checks for all 602 candidates, comprising 16,527 groups, 64,988 copies and 48,461 spacers. DNA verification used 179 archived contexts and 423 frozen-FASTA candidates, with no discrepancies. | Main Fig. 4; S9. | Native-DNA coordinate verification is complete according to the archived audit. This does not constitute a fresh 919-genome scan, establish availability of all production inputs or demonstrate biochemical ART function. |

**Scope of structural validation.** The reported structural evidence consists of automated comparisons; the J05 analysis did not include additional manual expert assessment. J05 supplies 2,144 completed US-align comparisons for 67 available CIFs against 32 fixed references within its 69-membership scope, retaining two historical technical failures. All 32 strict-mirror members, including the 18 strong and six suggestive members, are computationally covered; none is thereby marked as manually validated. The 69-row worksheet documents the comparison scope and does not provide completed manual expert assessments. Pairwise outputs and completion records are provided separately from the archived full-hit results. These automated comparisons do not replace expert structural review.

**Scope of functional inference.** The five 6.12-Gb computational candidate systems and the independent 54-Gb candidate panels were not tested for biochemical activity or native biological function in this study. Section 14 outlines proposed causal validation. Functional claims would require direct RNA/cDNA assays, appropriate catalytic controls and, where applicable, partner perturbation experiments. The strict-mirror 24/32 and novelty-only 12/32 classifications remain computational protein-level compatibility results. The proposed experiments remain prospective.

#### 19 Reference methods and software

The study uses ACNV [1], Prodigal [9], myRT/RVT-All [2], HMMER3 [3], MMseqs2 [4], Foldseek [6], Boltz-2 [5], PHROG [8], BLAST+ [10] and Logan [11]. US-align was executed as an orthogonal algorithm comparison for the 67 available CIFs and 32 references within the prespecified J05 scope [7]; this does not establish coverage of every structural panel or completed expert review. The UG27 reference annotations were obtained from the public Minerva example-locus release associated with Li et al. [12].

### 20 Exploratory E3A/E3B architecture-family analysis

The following E3A/E3B results were fixed on 7 October 2026. They provide exploratory evidence supporting architecture-first follow-up and are not a replacement for the later F3V2, 54-Gb or N1–N5 analyses.

#### 20.1 Broad families and heterogeneous topology

| Family | Frozen historical observation | Interpretive boundary |
|---|---|---|
| ARCH250_0010 | 7 genomes; 5 taxonomy-tail groups; 5/5 witness ORFs official-CDS supported; 57-aa witnesses; repeat upstream in all five; median gap 277 bp | A broad-coherent repeat–microprotein module; not a confirmed RT or new biological function |
| ARCH250_0001 | 48 candidates / 23 genomes / 12 taxonomy-tail groups; 35/38 exact-primary witness candidates with official support; upstream modal fraction 55.3% | Broad but mixed topology; potential multiple sub-architectures |
| ARCH250_0003 | 27 candidates / 20 genomes / 11 groups; 7/9 exact-primary witnesses official-supported; downstream modal fraction 55.6%; median gap 42 bp | Likewise heterogeneous: family size is not a single universal mechanism |

#### 20.2 Special-case controls

| Family | Historical observations | Open question |
|---|---|---|
| ARCH250_0025 | Three Lak-phage genomes; 3/3 exact-primary signatures; repeat upstream, median gap 9 bp; 0/3 matched to official CDS | Conserved missed short ORF versus gene-caller artefact. Translation evidence needed. |
| ARCH250_0036 | Two genomes, 2/2 official support, 265-aa hypothetical proteins; repeat downstream; 23-bp median gap; Cas4 context flag at 8.345 kb | Flagged CRISPR/defense neighborhood, not RT or retron evidence. |

A shared repeat core plus a nearby protein-family assignment indicates an architectural association within the screened universe. It does not imply homology novelty, mechanism, experimentally verified protein expression or independent cross-lineage biological replication.

#### 20.3 Computational evidence and validation boundaries

| Level | Computational evidence or experimental endpoint | Caveat |
|---|---|---|
| L0 | Raw-DNA ACNV plus generic positional recurrence | Motif and protein labels excluded from scoring |
| L1 | Five known ART-bearing loci dual-arm; ranks 1–4 | Historical known-locus characterization, not independent new ART truth |
| L2 | Non-array RT controls 0/3; UG27 near-neighbor specificity | Sequence recurrence is narrower than RT activity |
| L3 | Pilot 240 genomes: 383 candidates; broad 919 genomes: 3,674 mathematical regions | Region counts are not system counts |
| L4 | Historical D1/D4: 0/83 calibrated compact RT-fold support | No structural RT conclusion from uncalibrated weak hits |
| L5 | E1: 66 replicated families; E2A: 12 exact-primary high-specificity families | Replicated architecture, not verified function |
| L6 | 64/80 official CDS witness support; 63/64 hypothetical/uncharacterized | Official CDS supports coding context, not function |
| L7 | E3B lineage dereplication; lead ARCH250_0010 | Within-family source dependence remains relevant |
| L8 | Biochemical RT, RNA substrate, partner perturbation and native phenotype | Not performed in the present study |

#### 20.4 Biological interpretation and limitations

These historical results show why raw-DNA candidate nomination should not be confined to recoverable reverse-transcriptase identity. Some repeat-linked architectures recur with short, poorly characterized neighboring coding contexts. This is a computationally testable biological hypothesis; to elevate it, independent lineage-resolved conservation, expression/translation, structure/function and perturbation evidence are required.

Some Stage05E3B interpretations were revised in the subsequent F3V2 analyses. These exploratory observations do not independently establish new genomic architecture classes. The absence of calibrated RT-fold support remains a negative computational result under the stated thresholds.

### 21 Supplementary displays and native-DNA audit

#### 21.1 Supplementary Figures S1–S3

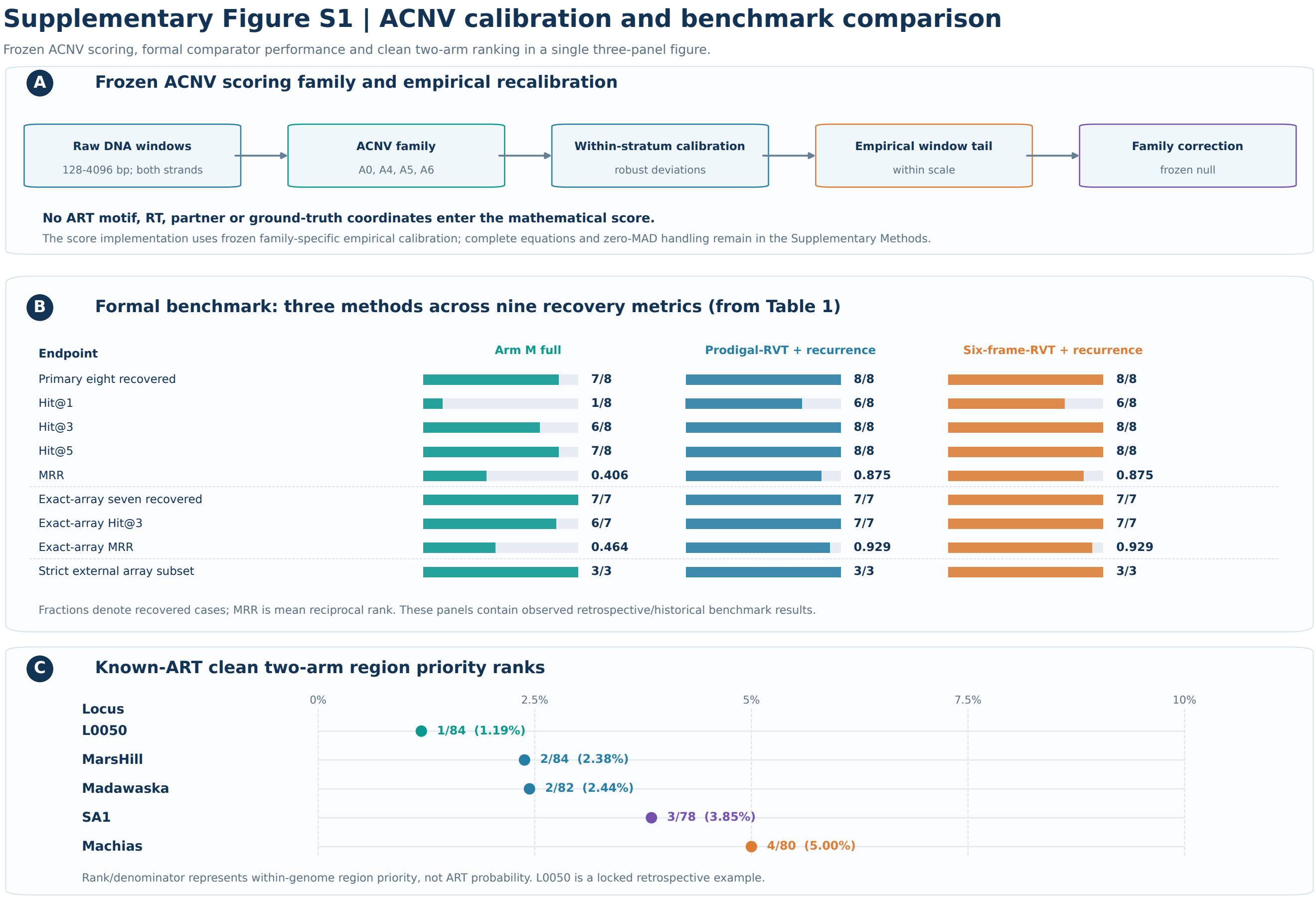


Supplementary Figure S1: ACNV calibration and benchmark comparison. Frozen ACNV scoring, formal comparator performance and clean two-arm ranking in a three-panel figure. Panel A summarizes frozen mathematical scoring and empirical calibration; panel B reports the formal benchmark (Table 1); panel C reports within-genome region priority ranks for five known ART-bearing loci.

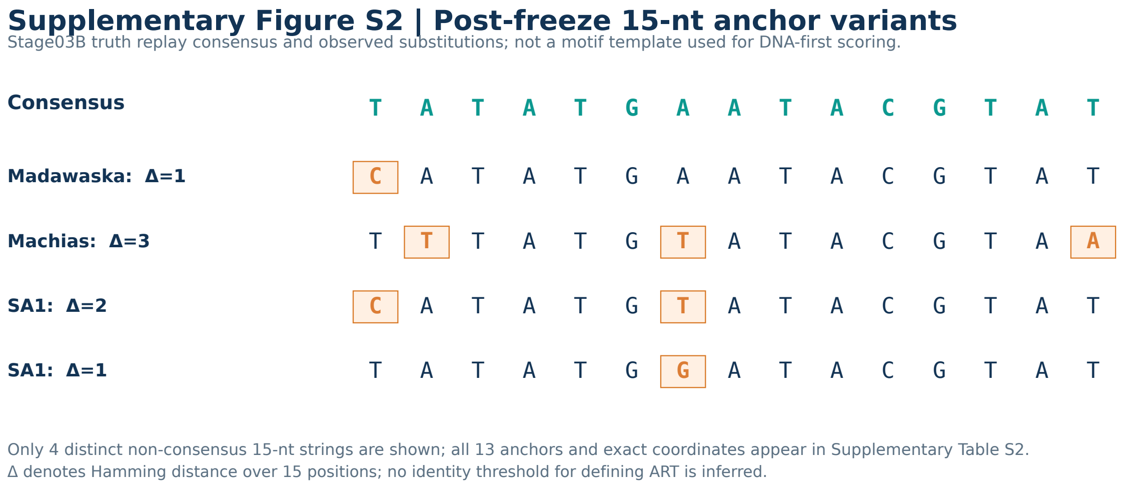


Supplementary Figure S2: Post-freeze 15-nt anchor variants. Stage03B truth-replay consensus and observed substitutions; these sequences were not used as motif templates for DNA-first scoring. Exact anchors and coordinates are provided in Supplementary Table S2.

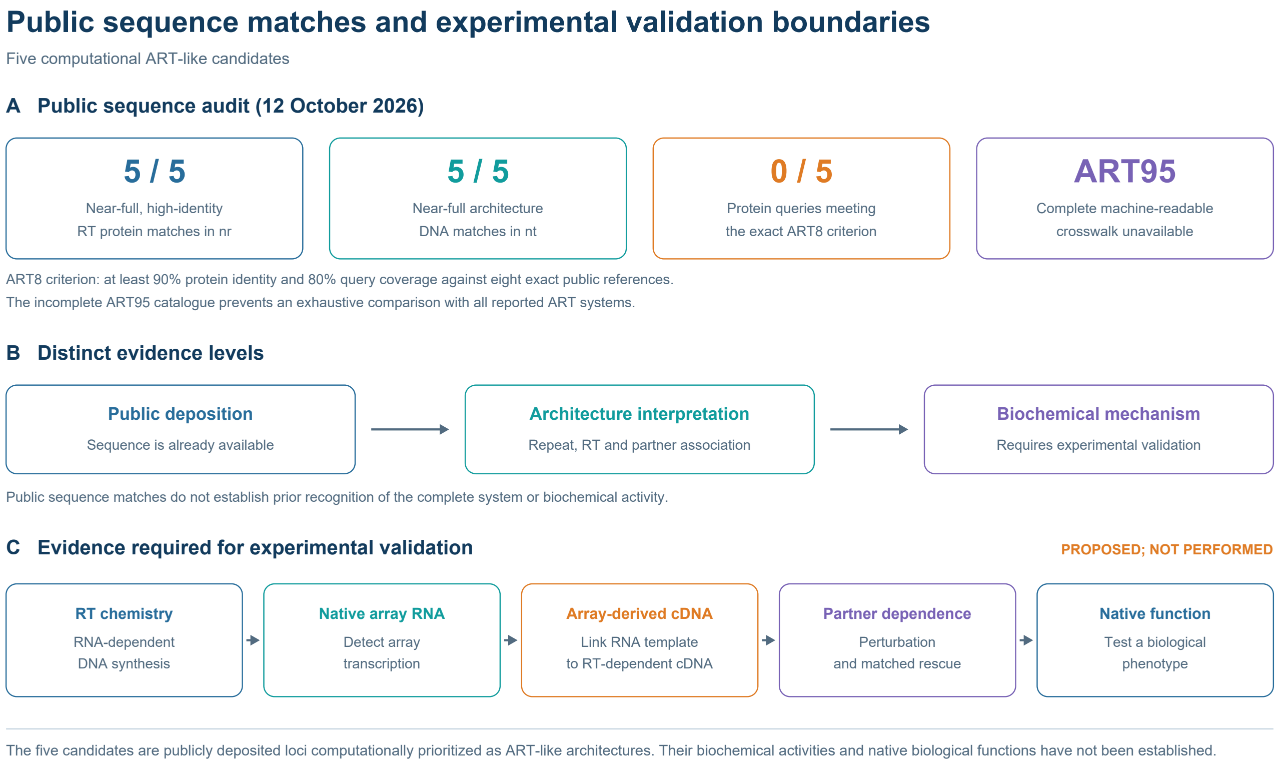


Supplementary Figure S3: Public sequence matches and experimental validation boundaries. **A,** The dated nr/nt audit records public protein and architecture-DNA matches for all five candidates; none of the five protein queries met the exact ART8 criterion of at least 90% identity and 80% query coverage against eight references. The incomplete machine-readable ART95 catalogue prevents an exhaustive crosswalk. **B,** Public deposition, architecture-level interpretation and biochemical mechanism are distinct evidence levels. **C,** The indicated assays are proposed requirements for validation and have not been performed.

#### 21.2 Supplementary Tables S1–S2

Supplementary Table S1: UG27 exact-unit dual-arm evaluation. Reference intervals were derived from published ncRNA annotations mapped after scoring outputs were fixed. Only A. putredinis has exact-unit dual-arm overlap (3/8); this does not independently establish biochemical function.

| System | Exact units | Dual units | Dual fraction | ACNV-only | Nearest dual gap (bp) |
|---|---|---|---|---|---|
| A. putredinis | 8 | 3 | 0.375 | 0 | 0 |
| B. coprophilus | 6 | 0 | 0 | 2 | 95,405 |
| Bacteroides sp. 9-1-42FAA | 6 | 0 | 0 | 2 | 10,968 |
| Bacteroides sp. 2-1-16 | 6 | 0 | 0 | 2 | 10,062 |
| B. hansenii | 2 | 0 | 0 | 2 | 26,086 |

Supplementary Table S2: Stage03B 15-nt motif variant catalogue. Replay consensus: TATATGAATACGTAT. Delta is Hamming distance across 15 bases. All anchors are post-freeze truth-replay annotations rather than motifs used for scoring.

| Genome | Accession | start0 | motif_15nt | Delta | Type |
|---|---|---|---|---|---|
| Madawaska | MW349129 | 213194 | CATATGAATACGTAT | 1 | variant |
| Madawaska | MW349129 | 213407 | CATATGAATACGTAT | 1 | variant |
| Madawaska | MW349129 | 213603 | TATATGAATACGTAT | 0 | canonical |
| Madawaska | MW349129 | 213815 | TATATGAATACGTAT | 0 | canonical |
| Madawaska | MW349129 | 213986 | TATATGAATACGTAT | 0 | canonical |
| Machias | MW349128 | 218595 | TTTATGTATACGTAA | 3 | variant |
| Machias | MW349128 | 218806 | TATATGAATACGTAT | 0 | canonical |
| Machias | MW349128 | 219002 | TATATGAATACGTAT | 0 | canonical |
| SA1 | MW218148.1 | 9596 | CATATGTATACGTAT | 2 | variant |
| SA1 | MW218148.1 | 9808 | TATATGAATACGTAT | 0 | canonical |
| SA1 | MW218148.1 | 9979 | TATATGAATACGTAT | 0 | canonical |
| SA1 | MW218148.1 | 10180 | TATATGAATACGTAT | 0 | canonical |
| SA1 | MW218148.1 | 10356 | TATATGGATACGTAT | 1 | variant |

#### 21.3 N2 native-DNA verification (24 October 2026)

The archived post-freeze N2 audit reports completed native-DNA checks of the repeat-group, copy and spacer coordinates for all 602 repeat-linked architecture candidates in the frozen 919-genome screen. The audit used original Stage05D1 repeat-group and copy records and original genome-native DNA context, without retuning the mathematical detector or repeating the full genome scan.

All 602 candidate IDs were accounted for: 262 exact-primary-repeat candidates and 340 approximate-repeat candidates. The retained qualified evidence includes 16,527 repeat groups, 64,988 repeat-copy records and 48,461 spacer intervals. The first 179 candidate contexts supported 20,887 copy and 15,593 spacer checks; the remaining 423 candidates supported 44,101 copy and 32,868 spacer checks using frozen genome FASTA files. Reported inconsistencies across the two batches: zero.

The 423-candidate second batch covered 236 genome accessions. Together, the two batches covered every member of the 12 E2A families (119 loci). The historical source report recorded successful native-sequence, interval and spacer checks on the original server; the large archived FASTA files were not included in the accompanying archive.

These archived results document the native-DNA coordinate verification summarized in P10; they do not establish availability or independent replay of all original production inputs. The coordinate checks do not establish biochemical ART activity in the 602 candidates or provide new independently adjudicated ART-positive ground truth. The source FASTA and provenance archives and the execution scripts remain on the project server and are subject to applicable publication and redistribution rights.

| Evidence | Status |
|---|---|
| Frozen candidate and copy accounting | Complete: 602/602 candidates; 64,988 copies |
| Native DNA coordinate check | Complete: 179 + 423 candidates |
| Independent ART biochemical activity | Not tested; experimental validation is required |
| Additional independently adjudicated ART benchmark | Not evaluated; N1 supplied zero independently validated positive loci |

*Source: original Stage05D1 / N2 native-423 and 179-candidate verification records, 24 October 2026.*
